# Gain-of-function cardiomyopathic mutations in RBM20 rewire splicing regulation and re-distribute ribonucleoprotein granules within processing bodies

**DOI:** 10.1101/2021.06.02.446820

**Authors:** Aidan M. Fenix, Yuichiro Miyaoka, Alessandro Bertero, Steven Blue, Matthew J. Spindler, Kenneth K. B. Tan, Juan Perez-Bermejo, Amanda H. Chan, Steven J. Mayer, Trieu Nguyen, Caitlin R. Russell, Paweena Lizarraga, Annie Truong, Po-Lin So, Aishwarya Kulkarni, Kashish Chetal, Shashank Sathe, Nathan J. Sniadecki, Gene W. Yeo, Charles E. Murry, Bruce R. Conklin, Nathan Salomonis

## Abstract

RNA binding motif protein 20 (RBM20) is a key regulator of alternative splicing in the heart, and its mutation leads to malignant dilated cardiomyopathy (DCM). To understand the mechanism of RBM20-associated DCM, we engineered isogenic human induced pluripotent stem cells (iPSCs) with heterozygous or homozygous DCM-associated missense mutations in RBM20 (R636S) as well as RBM20 knockout (KO) iPSCs. iPSC-derived engineered heart tissues made from these cell lines recapitulated contractile dysfunction of RBM20-associated DCM and revealed greater dysfunction with missense mutations than KO. Analysis of RBM20 RNA binding by eCLIP revealed a gain-of-function preference of mutant RBM20 for 3′ UTR sequences that are shared with amyotrophic lateral sclerosis (ALS) and processing-body associated RNA binding proteins (FUS, DDX6). Deep RNA sequencing revealed that the RBM20 R636S mutant has unique gene, splicing, polyadenylation and circular RNA defects that differ from RBM20 KO, impacting distinct cardiac signaling pathways. Splicing defects specific to KO or R636S mutations were supported by data from R636S gene-edited pig hearts and eCLIP. Super-resolution microscopy verified that mutant RBM20 maintains limited nuclear localization potential; rather, the mutant protein associates with cytoplasmic processing bodies (DDX6) under basal conditions, and with stress granules (G3BP1) following acute stress. Taken together, our results highlight a novel pathogenic mechanism in cardiac disease through splicing-dependent and -independent pathways that are likely to mediate differential contractile phenotypes and stress-associated heart pathology.

## INTRODUCTION

Dilated cardiomyopathy (DCM) is the most common indication for heart transplantation (McNally and Mestroni, 2017; Virani et al., 2021; Schultheiss et al., 2019). Recent insights into the genetics of DCM have revealed over fifty DCM causal genes that involve a wide variety of cellular processes and represent high priority targets for precision therapies (Schultheiss et al., 2019; McNally et al., 2013). These studies have expanded the scope of cardiomyopathy research well beyond cardiac energetics, conduction, or contractility (Jefferies and Towbin, 2010; Seidman and Seidman, 2011; McNally and Mestroni, 2017; Schultheiss et al., 2019; Norton et al., 2012). In particular, accumulating evidence implicates defects in RNA splicing and protein quality control in heart failure pathogenesis (Maatz et al., 2014; Rindler et al., 2017; Xu et al., 2005; Ding et al., 2004; Kong et al., 2010; Torrado et al., 2010). Indeed, DCM can be caused by mutation or dysregulation of multiple splicing factors (Anderson et al., 1995; Verma et al., 2013). Because the altered regulation of a single splicing factor can impact broad splicing regulatory networks, the effect of splicing factor mutations is multifaceted, leading to disruption of many cardiac signaling, transcriptional, and structural pathways.

The RNA binding motif protein 20 (RBM20) is a splicing regulator primarily expressed in heart and skeletal muscle, where it plays a central role in cardiac physiology. RBM20 directly binds to the primary RNA (pre-mRNAs) of many cardiomyopathy-associated genes where, by a process of exon exclusion, it ensures the proper production of adult protein isoforms (i.e., splice isoforms associated with cardiac maturation) (Maatz et al., 2014; Beraldi et al., 2014; Guo et al., 2012; Brauch et al., 2009; Wells et al., 2013). Autosomal dominant mutations in RBM20 account for up to 3% of DCM cases (Emig et al., 2010; Wells et al., 2013; Maatz et al., 2014). Further, RBM20 DCM is highly penetrant, is associated with life-threatening ventricular arrhythmias, and displays earlier age of onset than DCM associated with mutations in other proteins (e.g., laminA/C or Titin). RBM20 DCM missense mutations are enriched in a stretch of five amino acids (RSRSP, which contains the R636S mutation) in the arginine-serine-rich (RS) domain of the protein (Brauch et al., 2009; Emig et al., 2010). Surprisingly, nonsense mutations or missense mutations in other portions of the 1227-amino acid-long protein are relatively rare, suggesting that they are either undiagnosed due to a mild phenotype or not tolerated (Emig et al., 2010; Refaat et al., 2012).

Wild-type RBM20 has been shown to repress exon inclusion in key regulators of cardiac excitation-contraction coupling such as *TTN*, *CAMK2D* and *RYR2*, through binding to a UCUU consensus motif in adjacent introns (Maatz et al., 2014). Isolated cardiomyocytes (CMs) from rodents lacking RBM20 demonstrate prolonged action potentials and striking calcium handling defects including increased calcium transit amplitude, increased sarcoplasmic reticulum and diastolic calcium levels, and spontaneous calcium sparks (van den Hoogenhof et al., 2018). While loss of RBM20 expression leads to DCM, cardiac fibrosis, sudden death and arrhythmias in rodent models, few studies have focused on the precise role of mutant forms of the protein that correspond to human disease (van den Hoogenhof et al., 2018; Guo et al., 2012). Crucially, direct analysis of RBM20 RNA binding sites by CLIP (cross-linking immunoprecipitation) sequencing approaches has only been performed in wild-type (WT) rodent CMs and HEK293 cells (Maatz et al., 2014), thus whether mutant and wild-type RBM20 have distinct RNA binding preferences remains unclear.

Studies of the effect of pathogenic RBM20 mutants have spanned human, cell and animal models. Analyses of resected human DCM hearts from individuals carrying RBM20 mutations (R636S or S635A) have identified intriguing global differences in splice-isoform and circular RNA expression, which are associated with the regulation of cardiac contractility (Guo et al., 2012; Maatz et al., 2014). CM differentiation of human induced pluripotent stem cells (iPSCs) harboring pathological forms of RBM20 has identified calcium handling defects and splicing defects previously observed in rodent knockout (KO) models (Streckfuss-Bömeke et al., 2017; Wyles et al., 2016; van den Hoogenhof et al., 2018; Guo et al., 2012). In an impressive recent effort, pigs were engineered with either heterozygous (HTZ) or homozygous (HMZ) R636S mutations in RBM20 (Schneider et al., 2020). This model led to three crucial observations: 1) RBM20 R636S HMZ mutation leads to highly penetrant neonatal lethality due to heart failure; 2) mutant RBM20 co-localizes with stress granules in the CM cytoplasm after metabolic stress induced by sodium arsenate; and 3) RBM20 undergoes an apparent liquid-liquid phase separation. However, these studies raised intriguing questions regarding the role of RBM20 in pathogenesis, physiology, normal cardiac development and downstream RNA biology.

To address these questions, we used genome editing to generate an allelic series of RBM20 mutants, including one of the point mutations in the RS domain, R636S, as well as a RBM20 KO, in human iPSCs from a healthy subject. Normal splice patterns were differentially perturbed in RBM20 mutants as compared to KO, and genome-wide profiling of RBM20-RNA interactions revealed that mutant RBM20 showed a preference for the 3’ UTR of mRNAs. We observed co-localization of mutant RBM20 with cytoplasmic processing bodies, sites of mRNA storage and turnover, and with the canonical stress granule marker G3BP1 specifically following acute stress induction. Functional characterization of RBM20 R636S heterozygote 3D engineered heart tissues (EHTs) recapitulated aspects of DCM, providing a powerful model system. Collectively, our results indicate both splicing-dependent and splicing-independent mechanisms for RBM20 DCM pathogenesis, and suggest that RS domain mutant RBM20 has dominant-negative, gain-of-function properties.

## RESULTS

### Engineering iPSC-CMs with the RBM20 R636S mutation or RBM20 knockout

To model RBM20 cardiomyopathy mutants within iPSC-CMs, we selected a commonly occurring RS domain RBM20 missense mutation: R636S (DNA: C1906A) (Brauch et al., 2009). The heterozygous mutation (R636S HTZ) was introduced into iPSCs derived from a healthy male subject (WTC11) using transcription activator-like effector nucleases (TALENs) and a droplet digital PCR (ddPCR) based strategy (**Figs. 1A, B**) (Miyaoka et al., 2016, 2014). To create HMZ R636S mutant iPSCs, we retargeted the WT allele of the R636S HTZ iPSC line using an allele-specific sgRNA together with dual Cas9 nickases to create an allele with the R636S mutation and a silent mutation (T1905A) that distinguishes the newly created R636S plus silent mutation (SM) allele from the original WT and R636S alleles (R636S HMZ; **Fig. 1B** and **Figs. S1A-C**). To compare the R636S mutation with a loss-of-function mutation, we utilized our previously generated WTC11 iPSC line with an RBM20 HMZ 8-bp deletion (1917-1924del; RBM20 KO) (**Fig. 1C**) (Miyaoka et al., 2016). We confirmed that these cell lines maintained the normal male karyotype as well as loss of RBM20 protein in KO iPSC-CMs by immunofluorescent staining (**Extended Data Figs. 1D-F**) (Miyaoka et al., 2016). Importantly, we were able to generate iPSC-CMs from RBM20 mutants of high purity in all four cell lines (**Extended Data Fig. 1G**).

**Figure 1.**
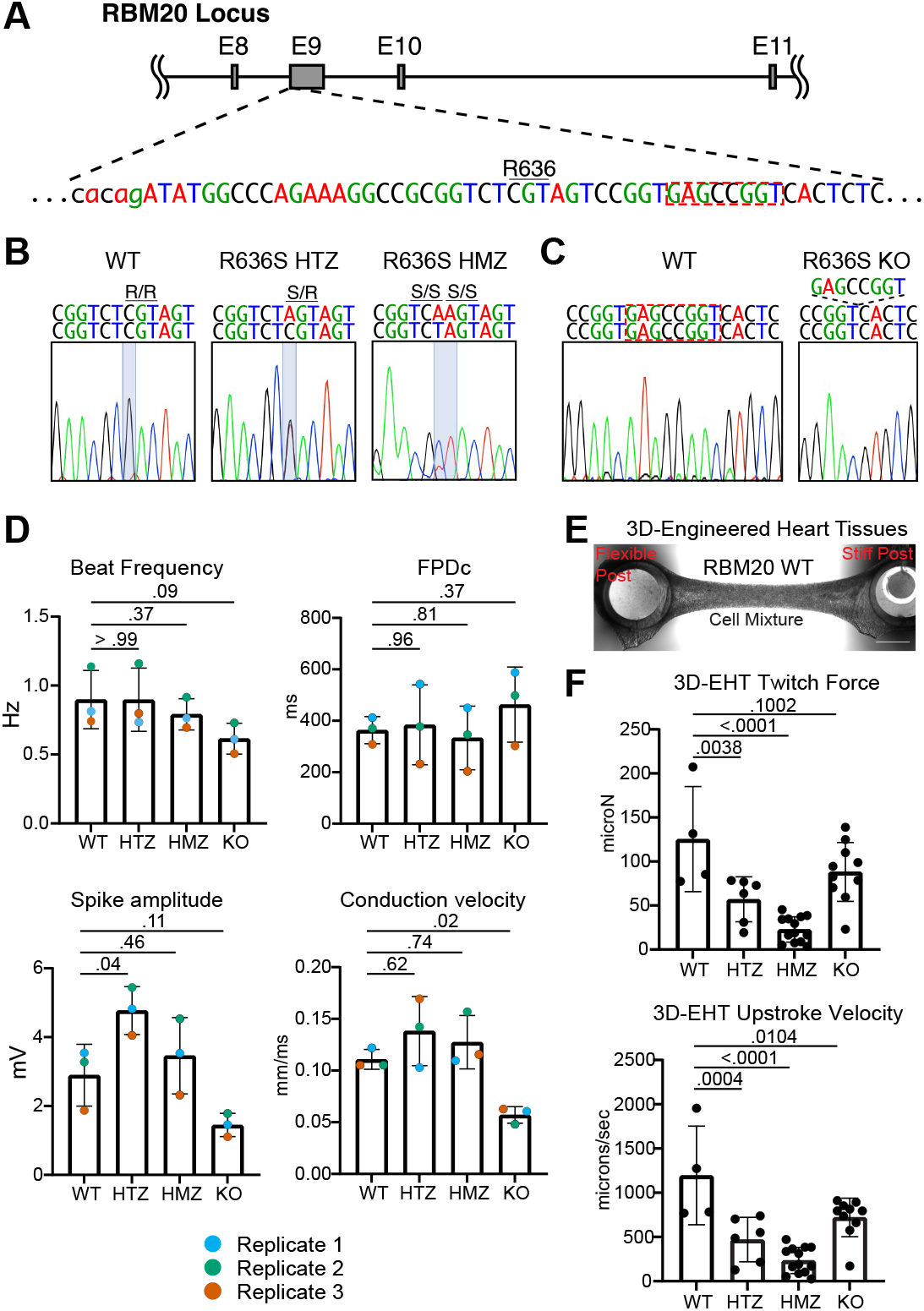
Generation and functional characterization of Mutant iPSC-CMs. A) Targeted genomic region of RBM20. The location of R636 and the 8 nucleotides deleted in 8-bp Del HMZ line are highlighted. B) Genomic sequences of the generated RBM20 HTZ and HMZ iPSC lines. C) Genomic sequence of the generated RBM20 8-bp Del HMZ line (KO). D) iPSC-CM electrophysiological parameters from Multi-electrode Array (MEA) recordings. Color-coded paired replicates represent a biologically independent average of 4 technical replicates. E) Representative 3D-engineered heart tissues (3D-EHT) from WT and RBM20 mutant cell lines. Phase-contrast microscopy, scale bar: 1 mm. The location of flexible and glass-stiffened silicone posts is indicated. F) Contractile metrics of 3D-EHTs. Dots represent individual 3D-EHTs from N = 2 for WT and N=3 for HTZ, HMZ, and KO 3D-EHTs independent biological replicates. Statistical significance was calculated using one-way ANOVA with Dunnett’s multiple comparisons test.

### R636S mutant iPSC-CMs exhibit electrophysiological and contractile abnormalities

Approximately 30% and 44% of RBM20 DCM patients display conduction system disorders and malignant ventricular arrhythmias, respectively. To test whether RBM20 mutant iPSC-CMs display electrophysiological aberrations, we used multi-electrode arrays (MEAs) to record their spontaneous field potential changes (**Fig. 1D** and **Extended Data Video 1**). While we observed some biological variability across different iPSC-CM batches, the trends between biological replicates were consistent (**Fig. 1D**). Field potential duration corrected for beat rate differences (FPDc), an *in vitro* surrogate of the QT interval and biomarker for arrhythmogenesis risk (Blinova et al., 2018), was not altered in RBM20 mutants. Beat rate was non-significantly reduced in RBM20 KO cells (p = 0.09), and RBM20 R636S HTZ iPSC-CMs displayed a statistically significant increase in spike amplitude, while RBM20 KO iPSC-CMs displayed a decrease in spike amplitude (**Fig. 1D**). In addition, conduction velocity was significantly decreased specifically in RBM20 KO iPSC-CMs (**Fig. 1D**).

The hallmark of RBM20 DCM is decreased cardiac contractility. To assess cardiac contractility in RBM20 mutant iPSC-CMs, we generated three-dimensional engineered heart tissues (3D-EHTs), an established model to promote iPSC-CM maturation closer to adult-like myocardium and perform more predictive disease modelling experiments (**Fig. 1E** and **Extended Data Videos 2, 3**) (Ronaldson-Bouchard et al., 2019; Campostrini et al., 2021; Karbassi et al., 2020; Leonard et al., 2018). Importantly, to ensure comparability across conditions we enriched hiPSC-CMs to >90% purity through metabolic selection with sodium lactate prior to 3D-EHT casting (**Extended Data Fig. 1G**). RBM20 R636S HTZ and HMZ 3D-EHTs displayed significantly decreased force generation and upstroke velocity, recapitulating aspects of DCM (**Fig. 1F**). Intriguingly, RBM20 KO 3D-EHTs demonstrated a less dramatic decrease than the RBM20 point mutations in both these parameters and were not significantly altered compared to WT controls (**Fig. 1F**). These data are in line with a recent report indicating that RBM20 KO mice have normal cardiac contractility (Ihara et al., 2020). Collectively, these electrophysiology and contractility analyses demonstrate distinct functional effects of RBM20 missense and nonsense mutation, suggesting that R636S may be a gain-of-function mutation.

### RBM20 is mis-localized in R636S mutant iPSC-CMs

To better understand the cellular mechanisms underlying the DCM-like phenotypes in R636S mutant HTZ and HMZ iPSC-CMs, we examined intracellular localization of the mutant protein. RBM20 is known to localize to the nucleus, where it forms discrete foci that exquisitely co-localize with the site of *TTN* transcription (Bertero et al., 2019). However, several groups have demonstrated that both RSRSP deletion mutants and DCM point mutations within the RSRSP stretch, including the R636S mutation, result in mis-localization of overexpressed (exogenous) RBM20 from the nucleus to the cytoplasm (Ihara et al., 2020; Schneider et al., 2020; Filippello et al., 2013; Murayama et al., 2018; Sun et al., 2020; Gaertner et al., 2020). Confocal analyses indicated that endogenous RBM20 in WT iPSC-CMs localized to the nucleus and was enriched in prominent foci (usually two per nucleus, as immature iPSC-CMs are generally diploid), while R636S HTZ and R636S HMZ iPSC-CMs displayed more numerous and smaller RBM20 puncta which appeared primarily cytoplasmic (**Fig. 2A**). To assess this aspect more robustly and exclude any potential artifact of maximum intensity image visualization of diffraction-limited microscopy, we turned to structured illumination microscopy (SIM), which affords a ~2x increase in both lateral and axial resolution compared to conventional techniques (e.g., confocal and wide-field; **Fig. 2B**) (Gustafsson, 2005, 2000; Heintzmann and Gustafsson, 2009). 3D axial projections of SIM data confirmed that RBM20 in R636S HTZ and HMZ iPSC-CMs was present in *both* the nucleus and cytoplasm, with a majority of RBM20 in R636S HMZ cells localized to the cytoplasm (**Fig. 2B, C**). These results indicate that contrary to previous reports, the exclusion of mutant RBM20 from the nucleus is not complete.

**Figure 2.**
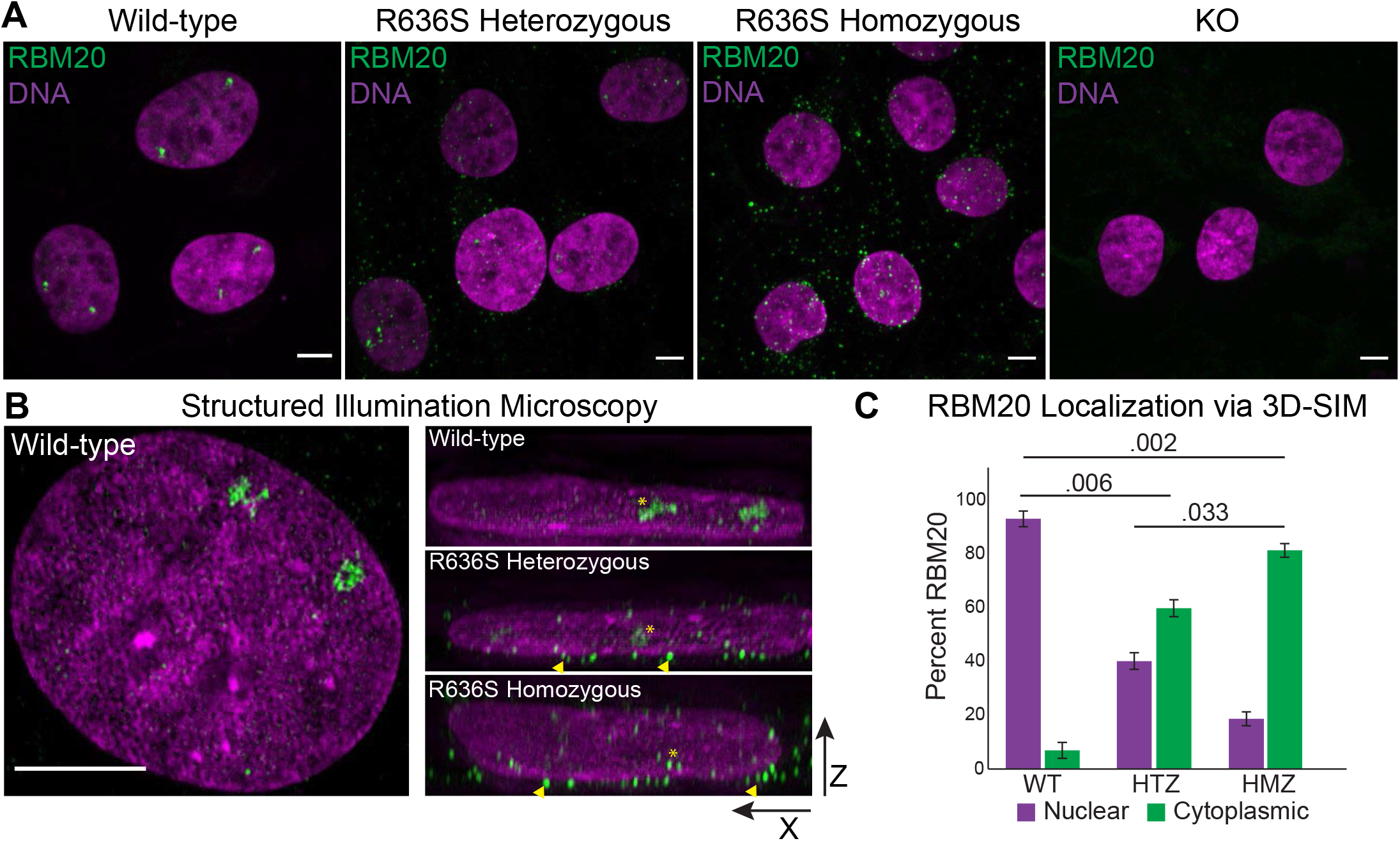
Intracellular RBM20 localization in RBM20 Mutant iPSC-CMs. A) Immunofluorescence for RBM20 in WT and RBM20 mutant iPSC-CMs. Spinning disk confocal, scale bar: 5 μm. B) Structured illumination microscopy of DNA (magenta) and RBM20 (green). Left WT image represents X-Y maximum intensity projection, right images represent X-Z 3D projections. Yellow arrowheads indicate perinuclear localization of RBM20 in mutant cells. Yellow stars indicate nuclear localization of RBM20. Scale bar: 5 μm. C) 3D localization analysis of RBM20 from 3D-SIM images. Magenta bars (left bar above genotype) indicate percent nuclear localization, green bars (right bar above genotype) indicate percent cytoplasmic localization. Data represents per-cell averages from N = 2 independent experiments.

### R636S and WT RBM20 mediate distinct mRNA interactions

To define the molecular mechanisms underlying R636S-associated defects leading to DCM, we performed an unbiased survey of direct targets for WT and mutant RBM20 binding using enhanced cross-linking immunoprecipitation sequencing (eCLIP) (**Fig. 3A** and **Extended Data Fig. 2A**) (Van Nostrand et al., 2016). RBM20 eCLIP of WT iPSC-CMs identified 1,240 reproducible peaks in biological replicates corresponding to 204 genes, including numerous well-established RBM20 splicing targets (e.g., *TTN, CAMK2D, RYR2, LDB3, LMO7, LRRFIP1, MLIP, OBSCN, SORBS2, TNN2*) (**Extended Data Table 1, 2**). Notably, these include prior presumed RBM20 direct targets not previously evidenced in rat heart HITS-CLIP (*CAMK2D*, *OBSCN*, *LDB3*) (Maatz et al., 2014). In addition, this analysis identified a number of novel RBM20 targets including ion channels (*CACNA1C, KCNIP4, KCNQ5, SLC8A1*), RNA binding proteins (*RBM20, FUS, QKI, FUBP3*), lncRNAs and other cardiac regulatory factors (e.g., *MYOCD*, *CTNNB1, HAND2, MYH6, SPATS2, GSE1, GNAI3, TPM1*). Consistent with the literature, WT RBM20 bound principally to intronic elements, and was most significantly associated with the established RBM20 consensus motif (UCUU) (**Fig. 3B, C**). In contrast, eCLIP of R636S HMZ iPSC-CMs identified 18,310 peaks in 2,794 genes. Most notably, more than 70% of RBM20 bound sequences within R636S HMZ iPSC-CMs were within the 3′ UTR of transcripts (**Fig. 3B, Extended Data Table 3**) and overlapped WT peaks in only 29 genes (e.g., *CACNA1C, TTN, GNAI3, QKI*). While the canonical RBM20 binding site remained the most enriched RNA recognition element (RRE) for R636S, secondary enrichments were found for *PTBP1* and *FUS* binding sites, RNA-binding proteins (RBPs) that are known to bind to the 3′ UTRs of transcripts in stress granules (**Fig. 3C**).

**Figure 3.**
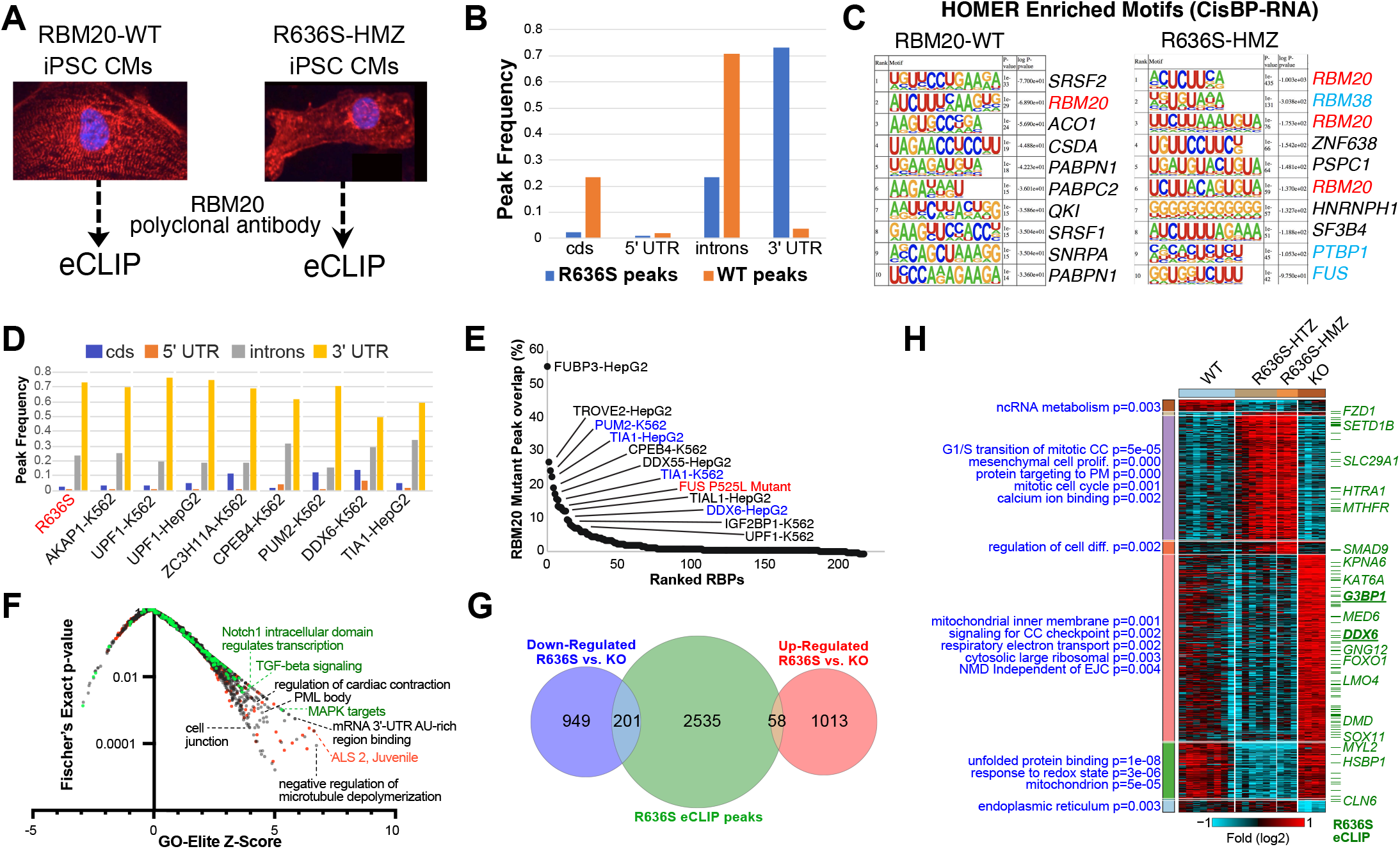
RBM20 mutant protein preferentially binds to the 3′ UTR of novel transcripts. A) Illustration of the eCLIP strategy for wild-type and R636S HMZ iPSC-CMs. B) Frequency of reproducible eCLIP peaks for WT and R636S HMZ iPSC-CMs within coding sequence (cds) exons, introns and UTR regions. C) HOMER de novo motif enrichment logos and enrichment p-values for the top ranked RNA-binding protein recognition elements defined from the CisBP-RNA database. RBM20-associated motifs are highlighted in red and RNA-stabilization associated factors in blue. D) Frequency bar chart of reproducible peaks in R636S HMZ eCLIP and the most similar eCLIP profiles from ENCODE (K562 or HepG2 cells), based on correlation to their frequency profiles. E) Overlap of R636S peaks in reproducible ENCODE eCLIP peaks, ranked by their percentage overlap. One previously described ALS eCLIP profile for the FUS-P525L mutation is included and highlighted in red, representing peaks only found in the FUS mutant compared to controls. Blue text indicates RBPs with statistically enriched RBM20 motifs-based on HOMER (Supplementary Fig. 4B). F) Gene-set enrichment with GO-Elite of genes associated with FUS-P525L and R636S overlapping peaks, for disease associated gene-sets (red, DisGeNET), aggregate pathways (green, ToppFun) and Gene Ontology (black). G) Overlap of genes up- or down-regulated by RNA-Seq in R636S iPSC-CMs (HTZ + HMZ) versus RBM20 deletion and R636S eCLIP peaks. H) Heatmap of all statistically ranked and organized (MarkerFinder algorithm) RBM20 R636S or deletion genes in iPSC-CMs by RNA-Seq. Statistically enriched gene-sets (GeneOntology + PathwayCommons) are indicated in blue with their associated Fisher’s Exact test p-value and R636S eCLIP peaks indicated by a green dash (selected genes shown).

Given its cytoplasmic localization and previous indications that the R636S mutant interacts with stress granules (Schneider et al., 2020), we hypothesized that RBM20 R6363S might act as an RNA stabilization factor. To test this hypothesis, we compared the transcript localization distribution of RBM20 R636S to all other RBP eCLIP profiles from ENCODE (ENCODE Project Consortium, 2012). Indeed, the R636S eCLIP was most similar to the binding pattern of well-established processing-body (P-body)/stress-granule associated RBPs including *PUM2, DDX6* and *TIA1*, all of which have eCLIP peaks enriched for the RBM20 binding site (**Fig. 3D, Extended Data Fig. 2B**). Direct comparison of R636S eCLIP peaks with all ENCODE eCLIP profiles resulted in a similar distribution of overlapping RBPs, including *IGF2BP1* and *TIAL1* (Fig. 3E). Interestingly, mutant RBM20 eCLIP binding sites also overlapped with those of a mutant form of FUS associated with ALS (P525L) (**Fig. 3E**) (De Santis et al., 2019). While mutant RBM20 eCLIP peaks were enriched in RNA-stabilization and cardiac contractile genes (**Extended Data Fig. 2C, D**), common peaks from both FUS-P525L and RBM20-R636S were also enriched in 3′ UTR AU-rich binding (e.g., *ELAV1, ME3D, CPE3*), ALS (e.g., *SETX, DCTN3, TRAK2*), PML body, cardiac contraction, and cell junction genes (**Fig. 3F, Extended Data Table 4, 5**). To assess the potential functional significance of these RBM20-R636S binding events, we performed deep RNA-Seq on WT (n = 8), R636S HTZ (n = 6), R636S HMZ (n = 3) and KO (n = 4) day 26 iPSC-CMs purified by sodium lactate treatment. Notably, R636S eCLIP peaks were relatively frequent (17%) among genes that were down-regulated in R636S versus RBM20 KO iPSC-CMs (**Fig. 3G** and **Extended Data Table 6**). Looking more broadly at global patterns of gene regulation identified striking differences between WT and KO or R636S iPSC-CMs, but only a few genes were consistently regulated in KO and R636S versus WT or between R636S HTZ and HMZ iPSC-CMs (**Fig. 3H, Extended Data Fig. 2E** and **Extended Data Table 7**). Among genes with the dominant pattern of up-regulation in the KO and coincident down-regulation in R636S iPSC-CMs, genes with R636S eCLIP peaks were abundant, including those for RBPs (DDX6 and the IGF2BP1 binding partner G3BP1) (**Fig. 3H**). Taken together, these data indicate that R636S RBM20 shares common binding sites with stress granule/P-body RBPs and with ALS mutant FUS, many of which are down-regulated only with R636S.

### Alternative splicing differences in R636S and RBM20 KO CMs impact distinct physiological pathways

Prior RBM20 KO studies in mice and rats, and patient RNA-Seq studies have identified a large number of RBM20-dependent splicing events implicated in DCM pathophysiology. To investigate the role of RBM20-mediated splicing in cardiac development, we first performed deep RNA-Seq of wild-type iPSCs at various stages of differentiation (**Extended Data Fig. 3, 4** and **Extended Data Table 8-13**). To detect changes in global splice patterns, we used a combination of analysis software (AltAnalyze and MultiPath-PSI; see Methods for details), allowing for an unbiased and quantitative comparison of known and novel splicing events between time points (Emig et al., 2010; Salomonis et al., 2010; Itskovich et al., 2020). In normal iPSC-CM differentiation, we observed an RBM20 splicing program induced in lock-step with an increase in *RBM20* gene expression (**Extended Data Fig. 4J**). These include well-established RBM20 target exons in *TTN* and *CAMK2D*. Notably, all novel cardiac differentiation splicing events tested (n=17) were readily verified by RT-PCR or targeted sequencing, indicating that our splicing analysis predictions are highly accurate (**Extended Data Fig. 4H, I** and **Extended Data Table 12**). Having confirmed the sensitivity of these algorithms and the emergence of an endogenous RBM20 splicing program during differentiation, we performed a global splicing analysis in gene-edited RBM20 iPSC-CMs. In R636S HTZ, we identified 116 splicing events that differed from those in WT iPSC-CMs using a conservative LIMMA-based analysis (**Fig. 4A; Extended Data Table 14; Methods**). These include several prior validated targets affected in DCM patients with the RBM20 S635A mutation, such as *TTN, CAMK2D, OBSCN, RYR2, IMMT,* and *TNNT2*) (**Fig. 4B-D**, and **Extended Data Fig. 5B**) (Guo et al., 2012). Several of these splicing events showed a clear dosage-dependent splicing pattern and overlap with RBM20 WT eCLIP peaks, suggesting that they are direct targets (e.g., *TTN, CDC14B, GSE1, RYR2, SPATS2*), while others had the same extent of splicing-deregulation with one or two alleles of R636S (e.g., *NEO1, IMMT, DMD*) (**Fig. 4B, D-E**). Approximately 80% of the RBM20 R636 HTZ alternative splicing events found were associated with cassette-exon splicing, in particular exon inclusion, consistent with prior reports (**Fig. 4A**) (Maatz et al., 2014). Notably, we found that >50% of these R636S-dependent splicing events were secondarily observed in independent RBM20 DCM mutant edited iPSC lines from our laboratory and others (**Extended Data Fig. 5A; Extended Data Table 15-18**). To assess the potential developmental regulation of these splicing events, we examined them in our WT iPSC-CM differentiation time-course. This analysis finds that R636S results in a developmental reversion of splicing for several well-described and novel RBM20 target exons (e.g., *CAMK2D, TTN, GSE1*) (**Extended Data Fig. 4E, F; Extended Data Fig. 5A**), along with others verified by RT-PCR (**Extended Data Fig. 5I**).

**Figure 4.**
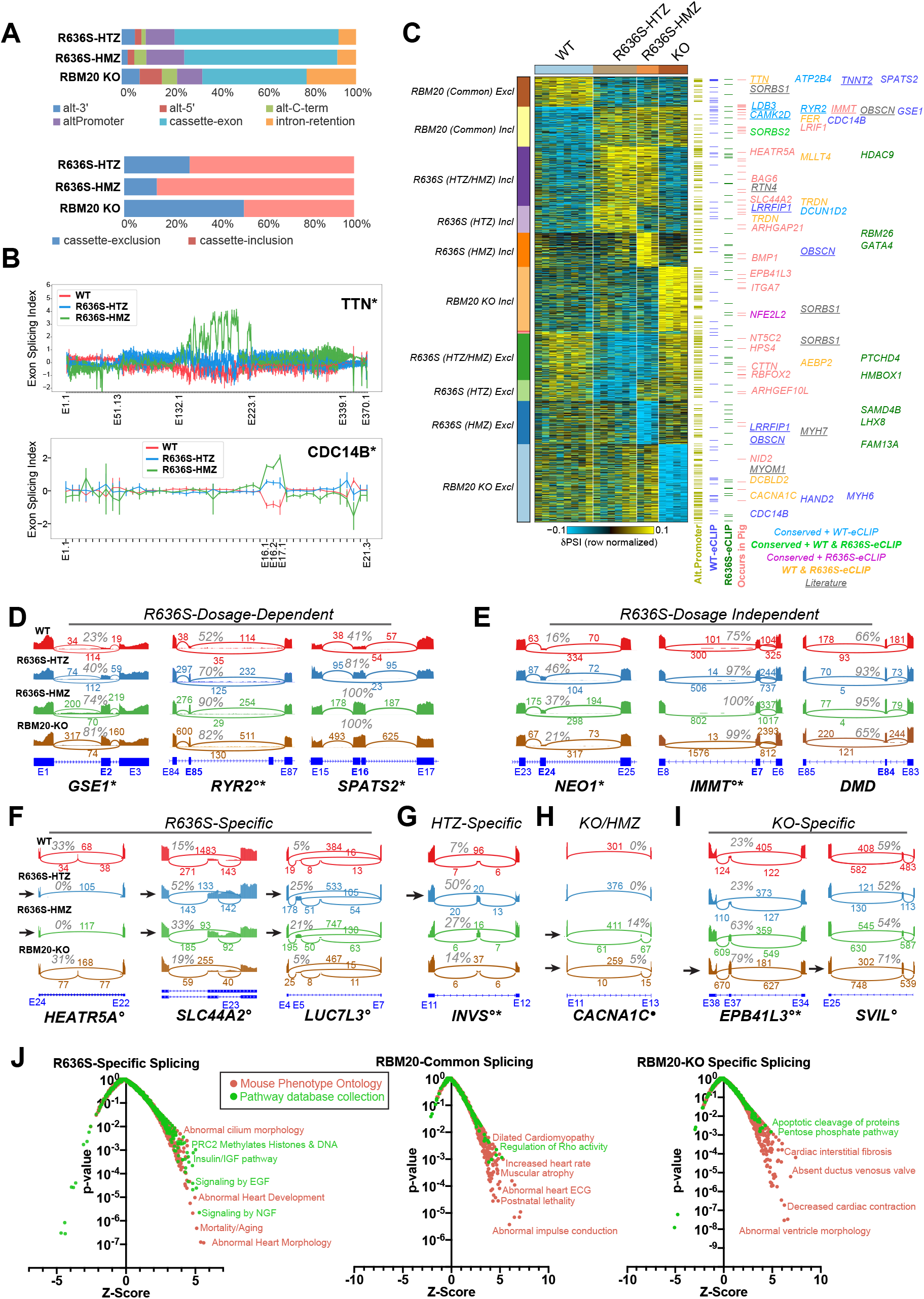
RBM20 mutation and knockout impact distinct pathways at the level of alternative splicing. A) Percentage of alternative splicing events considered differential (LIMMA corrected t-test p<0.1, FDR corrected and δPSI>0.1) for each of the iPSC-CM genotypes versus WT, separated by the predicted event-type (e.g., cassette-exon, intron retention) (top). Below, the percentage of splicing events associated with either cassetteexon inclusion or exclusion (skipping) are shown for each RBM20 genotype vs. WT. B) Gene-level visualization of exon-level relative expression levels (splicing-index) for two of example significant genes, *TTN* and CDC14B, predicted to be alternatively spliced in R636S HTZ and HMZ cells in a dosage-dependent manner. AltAnalyze exon identifiers are shown below. C) Heatmap of the predominant alternative splicing patterns (MarkerFinder) for all reasonably detected splicing events (eBayes t-test p<0.05, δPSI>0.1). The description of each pattern is displayed to the left of the heatmap (Incl = exon-inclusion, Excl = exon-exclusion associated splicing events). Splicing events with intronic eCLIP peaks in the same gene or that are also observed for the same exon-exon junctions as neonatal R636S porcine model (orthologous genome coordinates) are denoted to the right of the plot with examples listed. Underlined splicing-event genes indicate prior evidence of RBM20-dependent alternative splicing from rat KO studies. D-I) SashimiPlot genome visualization of RBM20-dependent splicing events observed in iPSC-CMs, associating with distinct patterns of regulation. Representative samples were selected. Specifically, R636S-allele dosage dependent splicing (D), dosage-independent R636S splicing (E), R636S but not KO dependent splicing (F), R636S-HTZ-specific events (G), R636S-HMZ-specific events, and RBM20-KO specific events (I) among a series of those visualized by SashimiPlot analysis (see Extended Data Fig. 5 and S6). Splice-junction read counts are denoted above each curved exon-exon junction line, along with the estimated percentage of exon-inclusion. J) Gene-set enrichment analysis (GO-Elite) of splicing-events segregated according to the MarkerFinder assigned patterns (panel C). Gene-sets correspond to either Mouse Phenotype Ontology or a collection of Pathway databases from ToppCell. * = verified splicing event patterns inferred from independently edited iPSC-CMs. ° = verified splicing event from R636S HTZ edited pig hearts. • = R636S eCLIP intron bound peak containing.

To identify predominant patterns of splicing in R636S mutation and RBM20 KO cells, we applied the same supervised-pattern analysis we applied to the gene expression data (**Fig. 4C; Extended Data Table 19; Methods**). This analysis revealed stark differences in the patterns of splicing compared to gene expression. Specifically, we observed splicing differences largely specific to the HTZ or HMZ R636S mutants, in addition to splicing differences uniquely shared by R636S HTZ and HMZ, those specific to RBM20 KO or shared among all R636S and RBM20 KO samples. The R636S/KO shared events frequently overlapped with WT eCLIP peaks and included the majority of prior validated RBM20 targets from human heart and rat RBM20 truncation mutant studies (Guo et al., 2012; Maatz et al., 2014). As these splicing events co-occurred in KO cells, they likely are the result of nuclear depletion of RBM20. Intriguingly, splicing-events specific to the R636S mutants or KO represented the dominant pattern of splicing and suggest that R636S does not simply phenocopy loss of RBM20 expression. Such R636S and KO-specific events included splicing regulators themselves (e.g., *LUC7L3*) as well as putative heart contractile disease mediators (*SLC44A2, SLCA81, SVIL*) (**Fig. 4F-I; Extended Data Fig. 5**). To determine whether such spliced exons are also present in an *in vivo* model of RBM20 DCM, we re-analyzed recent RNA-Seq from R636S gene-edited neonatal pigs, with HTZ and HMZ alleles (Schneider et al., 2020). This analysis verified splicing events within all observed splicing patterns that were conserved to pig at the level of exon-exon junctions (**Fig. 4C; Extended Data Fig. 6; Extended Data Table 20**). R636S-unique splicing patterns were further evidenced by R636S intronic eCLIP binding sites as well independent iPSC-CM RNA-Seq, with RBM20 missense and non-sense mutations (**Fig. 4C**). Gene-set enrichment analysis of RBM20 splicing events that fell into the three major patterns (commonly regulated, specific to KO, specific to R636S), highlighted distinct cardiac contractility, physiology and signaling pathways that represent potential areas for targeted exploration in the future, including ion channel-specific splicing differences found only in R636S (e.g., *TRPC3, CACNA1G, CACNA1D, CACNB3, KCNH2, KCNC4*) (**Fig. 4J** and **Extended Data Table 21**). In summary, we find that R646S and RBM20 KO result in largely distinct changes in splicing and that such human iPSC-CM events are conserved in a R636S pig model of DCM.

### Mutant and KO RBM20 alter circular RNA production and alternative polyadenylation

In addition to alternative splicing, recent studies suggest that circular RNAs, in particular those in *TTN*, may also contribute to RBM20-DCM pathology (Aufiero et al., 2018; Tijsen et al., 2021). Using an RNA-Seq protocol sensitive to circRNAs we found that R636S mutants produce a far greater number of circRNAs than RBM20 KO iPSC-CMs (**Extended Data Table 22**). Consistent with prior studies, the most frequently detected circRNAs were found in *TTN*, with relatively few differential circRNA exons/introns containing RBM20 eCLIP binding sites or overlapping with alternatively spliced-exons (**Extended Data Fig. 7A, B**).

While not previously implicated in RBM20 biology, we further asked whether alternative polyadenylation (APA) occurred in the setting of mutant or KO KBM20 from our RNA-Seq. Analysis of differential APA events using the QAPA algorithm highlighted ~1,200 events that were significantly associated with the dominant patterns of RBM20 mutant and KO regulation (**Extended Data Fig. 7C** and **Extended Data Table 23**). KO-induced APA events were enriched in mutant RBM20 eCLIP targets, which were predicted to mediate heart morphogenesis, the Golgi-network, cell-cycle, transcriptional regulation, cell-junctions, WNT and BMP signaling, as well as 3’UTR AU-rich binding (**Extended Data Fig. 7D, E** and **Extended Data Table 24**). These data suggest that APA may represent a novel form of RBM20-dependent gene regulation that alters the ability of RBM20 to associate with 3’UTR targets.

### R636S mutant RBM20 with co-localizes with cytoplasmic processing bodies

Based on our eCLIP comparative analyses, we predicted that mutant RBM20 may specifically localize within processing bodies (P-bodies), cytoplasmic ribonucleoprotein granules that exist in a liquid phase and may act to repress mRNA translation (Luo et al., 2018). Importantly, P-bodies are distinct from stress granules, which are transient protein-RNA ensembles that assemble in response to acute stress, and disassemble when the stimulus is removed (Protter and Parker, 2016; Wheeler et al., 2016), although current models conflict on the question of whether P-bodies and stress granules interact during acute stress (Hubstenberger et al., 2017; Chantarachot and Bailey-Serres, 2018; Luo et al., 2018). The protein DDX6 was highlighted as one of the most likely RBM20 interacting P-body candidates from our analyses, as it shares predicted binding sites and binding profiles with RBM20 R636S. High-magnification imaging revealed that a subset of cytoplasmic RBM20 in mutant iPSC-CMs co-localized with P-bodies as visualized with DDX6 (**Figs. 5A-C**). RBM20 co-localized with DDX6 to a greater degree in R636S HMZ than HTZ iPSC-CMs, and essentially no co-localization was observed in WT cells. Another P-body factor highlighted from our eCLIP analyses was IGF2BP1, which associates with the stress granule-promoting factor G3BP1. No stress granules were present in iPSC-CMs under basal culture conditions, in which G3BP1 displayed a diffuse cytoplasmic localization (**Fig. 5D**). However, following treatment with 1 mM sodium arsenate for 1 hour, iPSC-CMs rapidly assembled conspicuous stress granules (**Fig. 5D, E**). Under these acute stress conditions, mutant RBM20 co-localized with stress granules, but with WT protein did not (**Fig. 5E**). Hence, these data strongly support a model by which mutant RBM20 impacts alternative splicing through broad non-nuclear 3′ UTR binding and association with P-bodies, a mechanism that is reminiscent of neurological diseases associated with RNA-binding protein cytoplasmic aggregation (**Fig. 6**).

**Figure 5.**
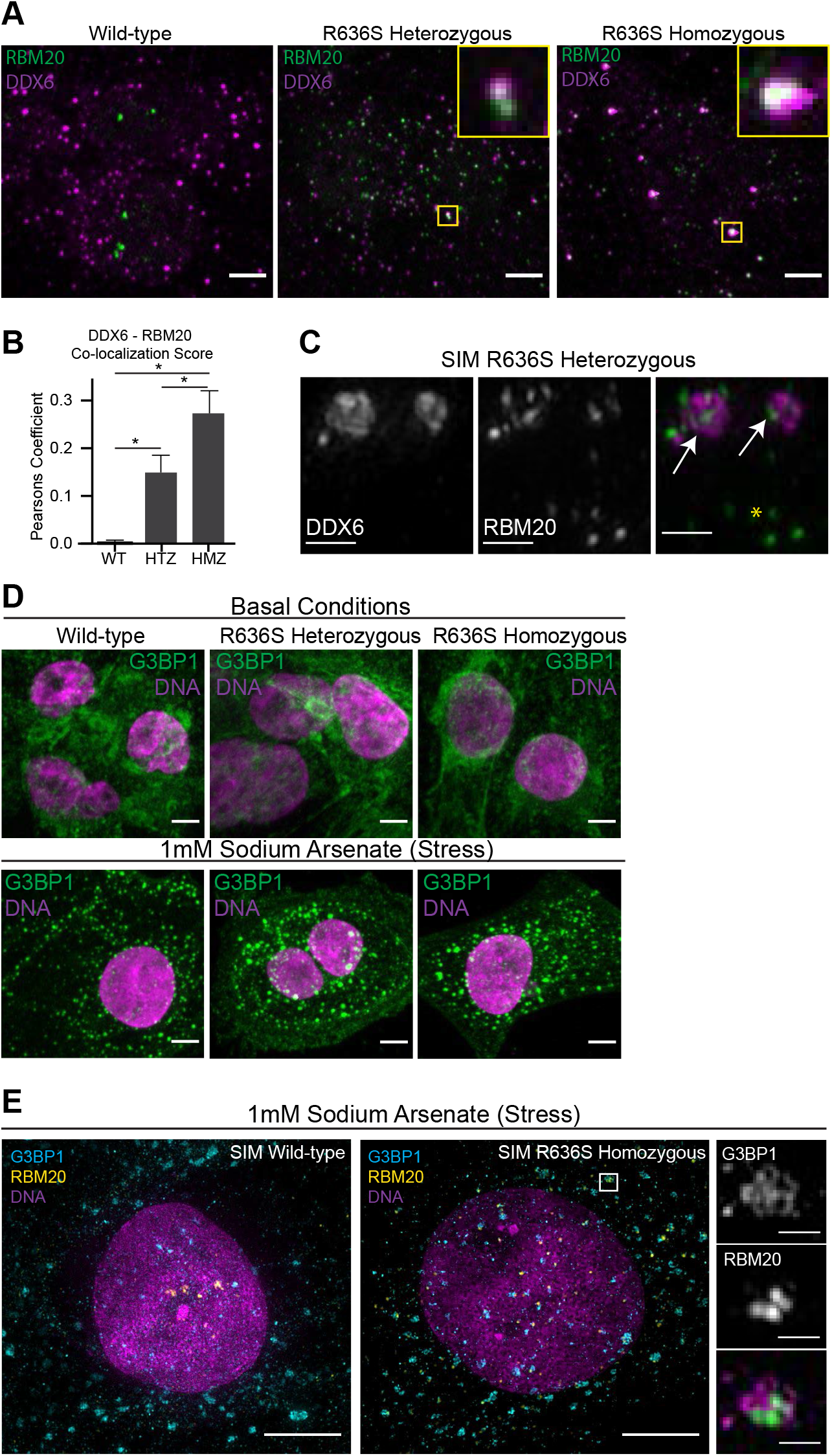
Intracellular localization of RBM20 mutant iPSC-CMs and association with P-bodies. A) Localization of RBM20 (green) and DDX6 (processing bodies, magenta). High-magnification inserts highlight co-localization). Spinning disk confocal, scale bar: 5 μm. B) DDX6/RBM20 co-localization is quantified via Pearson′s coefficient of co-localization. Data from N = 3 biological independent experiments. Statistical significance was calculated using one-way ANOVA with Dunnett’s multiple comparisons test. C) Structured Illumination microscopy of DDX6 (left), RBM20 (middle), and merged channels (DDX6: magenta, RBM20: green). White arrows indicate co-localization; yellow asterisk indicates RBM20 not associated with processing bodies. Scale bar: 0.5 μm. D) Assessment of stress granules via G3BP1 localization. iPSC-CMs were analyzed in basal conditions (e.g., normal media) and in response to stress (1 mM Sodium Arsenate treatment for 1 hour). Spinning disk confocal, scale bar: 5 μm. E) Structured illumination microscopy analysis of RBM20 (yellow) and G3BP1 stress granules (cyan) in sodium-arsenate treated iPSC-CMs. Scale bar: 5 μm.

**Figure 6.**
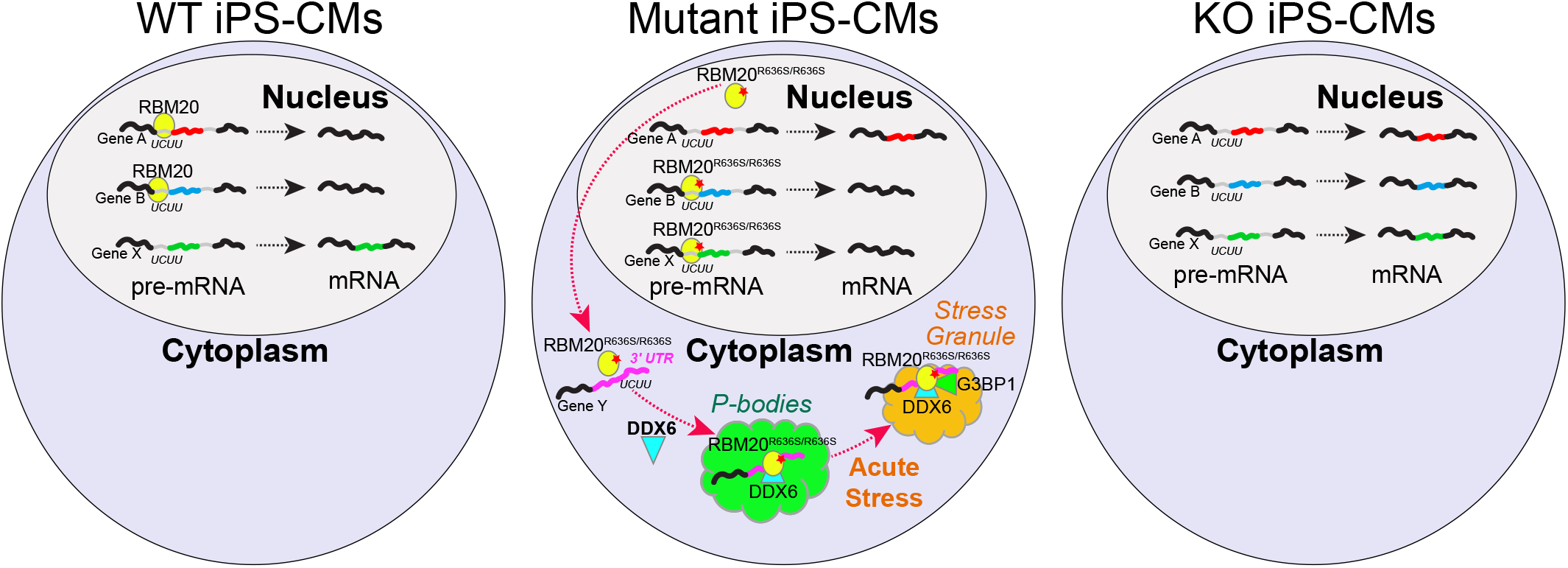
Model of mutant RBM20 differential splicing and P-body impacts in dilated cardiomyopathy. Proposed model for the impact of wild-type and mutant RBM20 on nuclear regulation of splicing based on RNA-Seq and eCLIP data as compared to cytoplasmic role of mutant RBM20 on P-body formation and 3’UTR association with mRNAs implicated in granule formation.

## DISCUSSION

Our understanding of the molecular and physiological effects of RBM20 mutations has remained elusive largely due to a lack of appropriate models that are not confounded by heart disease or patient genetics. Here, we applied precision targeting of an RBM20 patient genetic mutation in human iPSC-CMs as a model system to evaluate the impact of specific patient alleles and RBM20 KO. This controlled system allowed us to delineate the impact of RNA biogenesis on excitation-contraction coupling attributable to a single-variant substitution. We demonstrated distinct phenotypes via MEA between R636S mutant and RBM20 KO iPSC-CMs. Importantly, although previously reported (mouse, rat and pig) HTZ animal models of RBM20 DCM did not exhibit contractile defects, our RBM20 R636S 3D-EHTs (HTZ and HMZ) demonstrate a significant contractile phenotype compared to WT. Thus, this work establishes human iPSC-CM 3D-EHTs as a powerful and physiologically relevant model to study RBM20 DCM and therapeutic development.

Immunofluorescence experiments confirmed that the RBM20 R636S mutant protein mis-localizes to the cytoplasm. However, super-resolution microscopy revealed that a subset of mutant RBM20, even in R636S HMZ iPSC-CMs, maintains some capability to localize to the nucleus. Intriguingly, immunofluorescence showed that mutant RBM20 in mutant, but not WT iPSC-CMs co-localizes with P-bodies, cytoplasmic RNP granules present under basal conditions, that are sites of mRNA storage and turnover. During the preparation of this manuscript, the Schneider lab reported that mutant RBM20 co-localizes with stress granules in cells treated with sodium arsenate (Schneider et al., 2020). Stress granules are distinct from P-bodies, and are assembled in response to acute stress which inhibits translation (e.g., sodium arsenate, sorbitol, UV). Our data demonstrate that under basal culture conditions, WT and RBM20 mutant iPSC-CMs do not contain stress granules, but only assemble canonical stress granules in response to acute stress. Under these acute stress conditions, RBM20 in mutant, but not WT iPSC-CMs, does co-localize with stress granules. Schneider′s group also reported liquid-like behavior of mutant RBM20 (e.g., RBM20 potentially undergoes a “liquid-liquid phase separation”). As P-bodies have been demonstrated to represent a bona-fide liquid-liquid phase separated compartment (Hondele et al., 2019; You et al., 2020), association with P-bodies may drive this liquid-like behavior of mutant RBM20. Further investigation is warranted to better understand the functional consequences of the liquid-like behavior of cytoplasmic RBM20 on cellular physiology.

To understand the molecular consequences of mutant RBM20 in the nucleus and cytoplasm, we performed unbiased eCLIP profiling of wild-type and R636S iPSC-CMs. The results reveal a change in spliceosome-mediated target interactions in the setting of mutant RBM20 due to nuclear depletion of RBM20 and an altered bias towards consensus binding sites within the 3′ UTR of distinct transcripts. These 3′ UTR targets provide insight into the function of mutant RBM20, as many are shared with other RNA stabilization factors involved in pathogenic granule formation, including mutant FUS (Sabatelli et al., 2013). These data provide strong evidence that mutant RBM20 mimics previously described factors that mediate P-body and stress granule formation, and it may recruit additional factors to these bodies.

Importantly, our alternative splicing predictions further suggest a more complex model of the role of mutant RBM20 in cardiomyopathy pathogenesis. Specifically, although R636S RBM20 molecules are largely absent from the nucleus, we observed highly contrasting splicing signatures between KO and HMZ mutant cells. The importance of these mutation-specific splicing events, the targets of which include genes involved in cardiac development, and structural and contractile regulators, are supported by the finding of orthologous R636S-induced events in neonatal pigs and/or that exhibit overlapping mutant-specific eCLIP peaks. Additional observations from circular RNA analyses indicate that mutant RBM20 affects circular RNA production, supporting prior observations that such molecular events preferentially impact constitutive exons, while alternative polyadenylation analyses suggest a broader role RBM20 in post-transcriptional gene regulation (Aufiero et al., 2018). While the precise cause of and functional relevance of splicing differences in mutant and wild-type remain to be determined, these data suggest a more complex model in which loss of canonical RBM20 splicing leads to altered functional products involved in cardiac contractile regulation, potentially augmented by pathogenic circular RNAs, the emergence of new splice-forms in cardiac developmental and signaling genes, global changes in polyadenylation, and the degenerative accumulation of cytoplasmic granules impacting cardiomyocyte homeostasis.

It is intriguing to note the parallels between our observations with RBM20 and recent findings in neurodegeneration. Indeed, recent work has hypothesized cytoplasmic RBM20 may be similar to the cytoplasmic RNP granules associated with neurodegeneration (Schneider et al., 2020), such as TAU for Alzheimer′ s disease, Huntingtin for Huntington′ s disease, and FUS for amyotrophic lateral sclerosis (ALS) (Conlon and Manley, 2017; Maziuk et al., 2017). We add to this initial hypothesis by demonstrating that under basal conditions, mutant RBM20 binds to 3′ UTR regions of mRNA and co-localizes with P-bodies. Our study indicates that protein granules are a critical determinant factor for not only neurodegenerative disease but potentially more diverse diseases including cardiomyopathy. This is an intriguing notion, but further mechanistic studies are required to determine the functional consequences of the cytoplasmic localization and association of mutant RBM20 with P-bodies.

These analyses illustrate important differences in the splicing targets of mutant RBM20, with data from iPSC-CMs likely indicating that defects during early cardiogenesis could lead to remodeling of the fetal heart. While such molecular changes in iPSC-CMs may not recapitulate the gross-cardiac remodeling defects that impact heart function in patients with RBM20 mutations, we hypothesize that such splicing differences may predispose to later cardiovascular events. Importantly, these human cellular models provide the cardiovascular research community with powerful tools to aid in the development of new therapeutic targets for heart disease.

## ACKNOWLEDGMENTS

This work was supported by generous research grants from the National Heart, Lung, and Blood Institute (U01 HL099997, P01 HL089707, R01 HL130533, F32 HL156361-01, HL149734, R01 HL128362, R01 HL128368, R01 HL141570, R01 HL146868) to B.R.C, N.S, A.M.F., C.E.M., and N.J.S. National Institute of Diabetes and Digestive and Kidney (U54DK107979-05S1) to A.B. and C.E.M. National Science Foundation (NSF CMMI-1661730) to N.J.S. JSPS Grant-in-Aid for Young Scientists (A) [17H04993], NOVARTIS Research Grant, Mochida Memorial Foundation Research Grant, SENSHIN Medical Research Foundation Grant, Naito Foundation Research Grant, Uehara Memorial Foundation Research Grant, Uehara Memorial Foundation Research Fellowship, Gladstone-CIRM Fellowship to Y.M., and the A*STAR International Fellowship to K.K.B.T. Imaging was performed at the Gladstone Institutes’ Histology and Light Microscopy Core, Garvey Imaging Core at University of Washington, Biological Imaging Facility at University of Washington and iPSC work was carried out in the Gladstone Institutes’ Stem Cell core. We would like to thank Chi-Li Chiu for advice on quantitative image analysis, Samantha Bremner for advice and support on 3D-EHT experiments, and Deepak Srivastava and Shinya Yamanaka for their valuable advice on our data and the manuscript.

## DATA AVAILABILITY

The sequencing datasets have been deposited in GEO (GSE175886) and Synapse (https://www.synapse.org/#!Synapse:syn2582579), along with the processed data files.

## AUTHOR CONTRIBUTIONS

A.M.F., Y.M., A.B., S.S., S.J.M., M.J.S., K.K.B.T., J.P-B., P-L.S., G.W.Y., C.E.M., N.J.S., B.R.C., and N.S. designed the experiments. A.M.F. performed electrophysiological and contractile characterization of iPSC-CMs, and intracellular localization analysis of RBM20 with help from A.B. N.J.S. assisted with design and analysis of EHT experiments. Y.M., M.J.S., K.K.B.T., J.P-B., A.H.C., S.J.M., T.N., C.R.R., P.L., A.T., and S.S. generated genome edited iPSC-CMs and conducted RNA-Seq and eCLIP analyses. A.K., K.C., and N.S., performed computational analyses. A.B., P-L.S., G.W.Y., C.E.M., B.R.C., and N.S. supervised projects. A.M.F., Y.M., A.B., C.E.M, B.R.C., and N.S. wrote the manuscript with help from all authors.

## MATERIALS AND METHODS

### iPSC Culture

The UCSF Committee on human research #10-02521 approved the study protocol for iPSCs. The human iPSC lines used in this study were generated from a healthy male patient, WTC11 (Kreitzer et al., 2013; Miyaoka et al., 2014) using the episomal reprogramming method (Okita et al., 2011). Informed consent was obtained for this procedure. iPSCs were maintained on Matrigel (BD Biosciences) in Essential 8 medium (Life Technologies), which was exchanged every other day. To sparsely populated wells (i.e., passaged wells), we added 10 μM Y-27632, a Rho-associated kinase (ROCK) inhibitor (Millipore), to promote cell survival.

### RBM20 Mutant iPSC Line Generation

We applied our ddPCR and sib-selection based strategy to isolate genome-edited iPSC lines, as previously described (Miyaoka et al., 2016, 2014). We used TALENs targeting exon 9 of RBM20 (Addgene #108342 and #108343) to generate the RBM20 R636S Het iPSC line, which was then retargeted by CRISPR/Cas9 nickase (pX335; Addgene #42335) to convert the WT allele to the R636S with a S635S silent mutation allele. We used the Human Stem Cell Nucleofector Kit-1 and a Nucleofector 2b Device (Lonza) to transfect the plasmids and oligonucleotide donor DNA. For each transfection, 2 million cells were transduced with 3 μg of each TALEN vector or pX335 and 6 μg of an oligonucleotide donor DNA using program A-23. The transfected cells were plated into a Matrigel-coated 96-well plate using a multichannel micropipetter. The detailed information of the TALENs, gRNAs, and oligonucleotide donor DNA are summarized in **Supplemental Methods**. Karyotyping was performed by Cell Line Genetics.

The composition of the premixtures of allele-specific TaqMan probes and primers for ddPCR analysis was 5 μM of an allele-specific FAM or VIC TaqMan MGB probe (Thermo Fisher Scientific), 18 μM of a forward primer and 18 μM of a reverse primer (Integrated DNA Technology) in water. To detect point mutagenesis, we mixed the following reagents in 0.2 ml PCR 8-tube strips: 4 μl water, 12.5 μl 2× ddPCR Supermix for probes (Bio-Rad), 1.25 μl R636S+SM FAM probe and primer premixture, 0.625 μl WT VIC probe and primer premixture, 0.625 μl R636S FAM probe and primer premixture, and 5 μl (50–150 ng) genomic DNA solution (25 μl total volume). The conditions for droplet generation, thermal cycling, and data analysis for RBM20 R636S mutagenesis with the ddPCR system were described before (Miyaoka et al., 2014, 2016). As the R636S+SM FAM probe had a higher concentration than the WT FAM probe, the signal of the R636S+SM allele was distinguishable from that of the WT allele (**Extended Data Fig. 1**). Cell populations with a higher frequency of the R636S+SM allele and a lower frequency of the WT allele were enriched by sib-selection until the RBM20 R636S HMZ iPSC clone was isolated.

### iPSC-CM differentiation

For genomic analyses (Figs. 1, 3, and 4) The iPSC lines were differentiated into iPSC-CMs using the method that controls WNT signaling by small molecules (the GiWi protocol) (Lian et al., 2013). Briefly, iPSCs were seeded at 1.25-2.5×10^4^ cells/cm^2^ onto 12-well plates coated with 80 μg/μl growth factor-reduced Matrigel (BD Biosciences) in mTeSR1 supplemented with 10 μM Y-27632 (Millipore) for 24 h (day 3). mTeSR1 medium was changed daily for the next 2 days. On day 0, iPSCs were treated with 12 μM CHIR99021 (CHIR) (Tocris) in RPMI supplemented with B-27 (RPMI/B27) without insulin (Life Technologies) for exactly 24 h. On day 1, the culture medium was replaced with fresh RPMI/B27 without insulin and maintained for 48 h. On day 3, cells were treated with 5 μM IWP2 (Tocris) in RPMI/B27 without insulin and maintained for 48 h. On day 5, fresh RPMI/B27 without insulin was added to the cells, and on day 7, the medium was switched to RPMI/B27 with insulin. Afterward, fresh RPMI/B27 with insulin was added to the cells every 3 days. We used a metabolic selection protocol with glucose-free DMEM containing lactate to purify iPSC-CMs. Cells were replated on day 15-18 of differentiation, and then on day 20-22 media were replaced with DMEM (without glucose, with sodium pyruvate, Thermo Fisher Scientific) supplemented with Glutamax, non-essential amino acids, and buffered lactate (4 mM). Stock-buffered lactate solution was prepared by dissolving sodium L-lactate powder (Sigma-Aldrich) at 1 M concentration in 1 M HEPES solution. Lactate media were exchanged 2-3 times, with a total exposure of 48 hours for each treatment. After the final lactate treatment, media were changed to RPMI/B27 with insulin. The purified WT or RBM20 mutant iPSC-CMs at day 26 were harvested for RNA-Seq or cryopreserved for further analyses.

For electrophysiological, contractile, and immunofluorescence analyses, iPSCs were maintained on Matrigel (Fisher Scientific) and in mTesr Plus (StemCell Technologies) culture media, which was exchanged every other day. For iPSC-CM differentiation, iPSCs were plated in 12 well plates (Fisher Scientific) coated with Matrigel in mTesr Plus supplemented with 10 μM Y-27632 (StemCell Technologies) for 24 hours. On day −1, iPSCs were treated with 1 μM CHIR-99021 (Cayman Chemical) to prime cells for differentiation. On day 0, iPSCs were treated with RPMI-1640 with glutamine supplemented with 500 μg/ml BSA, 213 μg/ml ascorbic acid (RBA media; Millipore Sigma) and 5 μM CHIR-99021 for 48 hours. On day 2, cells were treated with RBA media supplemented with 2 μM WNT-C59 (Selleck Chemicals). On day 4, cells were treated with RBA. Starting on day 6, iPSC-CMs were maintained in RPMI + B27 (ThermoFisher) with media exchanges every 2 days. On day 13, iPSC-CMs were pre-conditioned with a 45-minute heat shock at 42°C (pre-warmed 42°C media was exchanged prior to incubation) to enhance survival of cryopreservation and thawing process. On day 14, iPSC-CMs were frozen in batches (>5 million cells) in CryoStor media (1 million cells/100 μLs CryoStor; Millipore Sigma) and placed in liquid nitrogen for disease modelling experiments. To passage cells for freezing, cells were washed with 1x DPBS (no magnesium, no calcium; ThermoFisher), and treated with 10x TrypLE (ThermoFisher) for 10 – 20 minutes at 37 °C. Cells were dissociated with mild trituration and passed through a 100 μM strainer (Fisher Scientific) to remove any large clumps (which negatively impact survival in our hands).

Frozen iPSC-CM stocks were thawed onto 6-well plates coated with Matrigel at 2.5 – 3 million cells/well in RPMI- B27 supplemented with 10 μM Y-27632 and 10% FBS. 24 hours post thaw, media was exchanged for RPMI-B27. After 48 hrs recovery, iPSC-CMs were purified in no glucose DMEM + 4 mM sodium lactate to select for iPSC-CMs for a total of 4 days (media was exchanged at day 2 of treatment). After 4 days, media was exchanged with RPMI-B27 and iPSC-CMs were allowed to recover for one day before re-platting (as above) for subsequent studies at day 21 post-differentiation.

### Flow-cytometry

Day 15 cells were dissociated to single cells using 0.2 mL of 0.25% Trypsin (UCSF Cell Culture Facility) per well of a 24-well plate with incubation at 37°C. Cells were resuspended in 0.8 mL/well EB medium (Knockout DMEM supplemented with 20% FBS (HyClone), Glutamax (Gibco), Non-essential Amino Acids (UCSF Cell Culture Facility), and 0.1 mM beta-Mercaptoethanol (Sigma)). 200 μl (about 200,000) cells were pelleted and fixed in 4% paraformaldehyde at room temperature for 15 minutes. Cells were permeabilized in FACS Buffer (DPBS without calcium and magnesium, 4% FBS, and 2 mM EDTA (Gibco)) with 0.5% (w/v) saponin (Sigma). Cells were stained with 100 μl of 0.002 μg/μl (1:100) Mouse anti-human cardiac Troponin T (cTnT) primary antibody (Thermo, MS-295-P) in FACS Buffer with saponin at room temperature for 30 minutes and washed. Cells were then stained with 100 μl of 0.01 μg/μl (1:200) Alexa Fluor 488 goat anti-mouse IgG secondary antibody (Invitrogen, A-11029) in FACS Buffer with saponin at room temperature for 30 minutes and washed. Finally, cells were stained with 100 μl of 0.01 μg/μl (1:1000) Hoechst 33342 (Molecular Probes) in FACS buffer at room temperature for 5 minutes and passed through a 0.4 μm filter (Millipore) to remove cell clumps. Data were collected using the MACSQuant VYB flow cytometer (Miltenyi Biotec) and analyzed using FlowJo.

### RNA-sequencing

Total RNA was purified from iPSC-CMs by using TRIzol and PureLink RNA Mini Kit (Thermo Fisher Scientific) according to the manufacturer’s instructions. The RNA-Seq libraries were prepared from the purified RNA by using TruSeq RNA Library Prep Kit v2 (Illumina) for gene and splicing analyses or the RiboMinus for Ribosomal RNA Depletion protocol (Thermofisher) for circular RNA splicing analyses. Paired-end RNA-Seq was performed with HiSeq (Illumina). An average of ~90 million reads were obtained from this iPSC RNA-Seq data. RNA-Seq for WTB iPS differentiation was performed as biological replicates (n=2), whereas the number of samples processed for WTC iPSC-CMs verified from 3-7 replicate samples per genotype.

### eCLIP

RBM20 WT and RM20-R636S direct transcript binding sites were determined using the same RBM20 polyclonal antibody (ThermoFisher, Cat#PA5-58068) on WT or R636S HMZ WTC iPS CMs using the eCLIP protocol as previously described (Van Nostrand et al., 2016). RNA-libraries were sequenced at a depth of 30-40 million reads for each replicate eCLIP sample, on an Illumina HiSeq 2500 (single-end reads). Reproducible RBM20 peaks (hg19) obtained from replicate WT and R636S HMZ iPSC-CMs compared to size-matched input controls, were used for all down-stream analyses. eCLIP peaks (hg19) from all prior generated ENCODE RNA-binding proteins (K562 and HEPG2 cell lines) were obtained from https://www.encodeproject.org/eclip/. Additionally, peaks associated with the ALS mutation in the gene FUS (P525L) were aggregated within a 200nt window using the software bedtools (merge function), only found in the mutant and not the FLAG tagged eCLIP, following liftover from hg38 to hg19 (GSE118347). To identify RNA recognition elements (RRE) associated with known RNA-Binding Proteins, we analyzed reproducible RBM20 peaks using the software HOMER using de novo motif discovery with the CisBP-RNA database (Ray et al., 2013; Heinz et al., 2010). All eCLIP peaks were annotated according to the gene intervals they correspond to (e.g., exon, introns, 3′ ends, 3′ UTR, distal or proximal regions) using predictions produced from clipper (https://github.com/YeoLab/clipper/). eCLIP peaks overlapping between different RBPs were identified using the bedtools intersect -wb option.

### RNA-Seq Analysis

RNA-Seq FASTQ files were aligned to the human hg19 reference genome and transcriptome using the software STAR. STAR produced BAM files were further processed in AltAnalyze (version 2.1.4) to obtain gene expression (RPKM values) and splicing estimates (PSI). Gene expression differences across the 11 CM differentiation time-points were calculated as pairwise comparisons to day 0 and to the prior time-point, for gene-set enrichment analysis and to restrict genes for pattern-based analyses (MarkerFinder algorithm in AltAnalyze) using an empirical Bayes moderated t-test p<0.05 (FDR corrected) and at least a 2-fold difference with a minimum RPKM value of 1 in either of the two groups compared. To accurately detect diverse alternative splicing events in iPSC-CM RNA-Seq, we used a recently described Percent Spliced-In splicing approach, called MultiPath-PSI, capable of accurate detection of known and novel exonic and intronic splicing events, as well as alternative promoter-associated exons (Itskovich et al., 2020; Muench et al., 2018; Rindler et al., 2017). Alternative polyadenylation was predicted and quantified using the software QAPA with default options to produce alternative isoform ratios for each 3′ UTR isoform, normalized to the total 3′ UTR form for each gene (Ha et al., 2018).

Application of MultiPath-PSI to all evaluated mutant and matched human WT samples identified 44,911 detected splicing events, corresponding to 16,207 unique splicing events. Prior differential splicing analysis, empirically observed gene expression associated differentiation effects were evaluated using unsupervised subtype detection analysis with the software Iterative Clustering and Guide-gene Selection (ICGS) (Olsson et al., 2016). Since the identified transcriptomic differences among independent iPSC-CM differentiations couldn′t be associated with iPSC genetic background, we surmise they are likely due to differentiation associated cardiac maturity. To account for these empirically observed effects, we applied a mixed effects linear model to account for these differentiation effects. For this model, each pair of differentiation-associated transcriptional effects was evaluated using LIMMA using the lmfit function to account for these effects in addition to cell line genetics in the linear model. See https://www.synapse.org/#!Synapse:syn25835429 for details.

For the differential splicing analysis, events were filtered for event detection in 75% of the samples in each biological group along with a change in PSI between the two groups greater than 10% (dPSI > 0.1). The supervised splicing pattern analysis was performed on unique (splicing-junction clusterID) identified differentially spliced events using the R packages LIMMA to obtain a two-tailed empirical Bayes moderated t-test p-value for the following models: 1) R636S HTZ-specific splicing, 2) R636S HMZ-specific splicing, 3) equivalent differential splicing in R636S HTZ and HMZ, and 4) additive induction in the HTZ to HMZ R636S. To identify splicing events associated with the predominant genetically-defined patterns, we applied the MarkerFinder algorithm, iterated over multiple sample groupings (all primary groups and aggregated groups of mutations - e.g., R636S-HTZ plus R636S-HMZ), for empirical Bayes moderated t-test significant events (p<0.05). This algorithm was also used for gene expression and alternative polyadenylation analyses, to define the major RBM20-dependent patterns (iterativeMarkerFinder function from https://git.io/Jt9je). Alternative splicing events from independently edited WTB iPSC-CMs with R636S HTZ and controls were quantified as above by TruSeq RNA-Seq, followed by AltAnalyze and MultiPath-PSI (δSPI>0.1). Similarly, RNA-Seq FASTQ files from a prior study were downloaded from PRJNA579336, to identify differential splicing results associated with missense or nonsense HMZ RBM20 mutations and controls, using AltAnalyze and MultiPath-PSI (δSPI>0.1 and empirical Bayes moderated t-test p<0.05) (Briganti et al., 2020). Gene-set enrichment analyses were performed with the software GO-Elite in AltAnalyze (Zambon et al., 2012).

To identify, quantify and annotate circular RNAs from back-splice junctions we first aligned the RNA-Seq results using TopHat2 on the RiboMinus depleted paired-end RNA-Seq FASTQ files, via the CIRCexplorer2 pipeline, on each individual sample (default options). This workflow identified far greater putative circRNAs than using the same pipeline with STAR 2.4.0. For differential circRNA analysis we ran RUVSEQ from the CSBB pipeline (https://github.com/csbbcompbio/CSBB-v3.0) on the back-splice junction counts for each sample for all relevant comparison groups. This workflow applies the software EdgeR to perform a differential expression analysis using a generalized linear model approach using the upper quartile normalization. All together, we identified circRNAs for 448 genes, with evidence of at least two reads per circRNA. circRNAs with a non-adjusted EdgeR p<0.1 were considered differentially expressed.

To determine genomic coordinate overlaps between: a) eCLIP peaks and alternative splicing events, b) eCLIP peaks and alternatively expressed circRNAs or c) alternative splicing events and alternatively expressed circRNAs, we created a multi-use script that leverages transcriptomic feature genomics coordinates and their overlap in AltAnalyze (https://git.io/JtHfa). For alternative splicing the MultiPath-PSI reported alternative exon and flanking intron genomic positions were directly intersected against eCLIP peaks. Similarly, the regulated exon(s) and flanking introns were considered for overlap for circRNAs.

### Quantitative PCR Analysis

Gene expression analysis was performed on the total RNA isolated during iPS cell differentiation into CMs. cDNA was generated from 1 μg of TurboDNAse-treated (Ambion) total RNA with the SuperScript III First Strand Synthesis kit and random hexamers (Invitrogen) as described by the manufacturer. Expression was assessed using TaqMan probesets run on the 7900HT real-time thermocycler (Applied Biosystems). Samples were assayed in technical triplicate, normalized to GAPDH or UBC, and relative expression was calculated with the day of highest expression set to 100%.

### Automated Primer Design and RT-PCR Analysis of ASEs

To validate alternatively spliced exons, the primer design program Primer3 (Rozen and Skaletsky, 2000) was integrated with AltAnalyze (identifyPCRregions function in the EnsemblImport module). As sequence input the alternative spliced exon, junctions and associated isoforms were used to identify the target region in the inclusion isoform (included/excluded exon), and suitable upstream and downstream exon sequences shared by the two isoforms. RT-PCR was conducted using these primers (**Extended Data Table 12**), random hexamer generated cDNA (described above), and Platinum Taq DNA Polymerase High Fidelity (Life Technologies) as described by the manufacturer.

### Engineered Heart Tissue Casting

3-dimensional engineered heart tissues (3D-EHTs) were cast and characterized as previously described (Leonard et al., 2018). Polydimethylsiloxane (PDMS) posts were fabricated by pouring uncured PDMS (Sylgard 184 mixed at a 1:10 curing agent to base ratio) into a custom acrylic mold (Limited Productions Inc., Bellevue, WA; design available upon request). Glass capillary tubes (1.1 mm in diameter; Drummond) were cut to length and inserted into the holes on one side of the mold before curing to render one post in each pair rigid. Post racks were baked overnight at 65°C before being peeled from the molds. Racks consisted of six pairs of posts that were evenly spaced to fit along one row of a standard 24-well plate. Fabricated posts were 12.5 mm long and 1.5 mm in diameter with a cap structure (2.0 mm in diameter for the topmost 0.5 mm) to aid in the attachment of 3D-EHTs. The center-to-center post spacing (corresponding to pre-compacted 3D-EHT length) was 8 mm. Prior to casting 3D-EHTs, all 3D-printed parts and PDMS posts were sterilized in a UVO Cleaner (342; Jetlight) for 7 min, submerged in 70% ethanol, and rinsed with sterile deionized water. Rectangular 2% wt/vol agarose/PBS casting troughs (12 mm in length, 4 mm in width, and ∼4 mm in depth) were generated in the bottom of 24-well plates by using custom 3D-printed spacers (12 mm × 4 mm in cross-section and 13 mm long) as negative molds. PDMS post racks were positioned upside down with one rigid-flexible post pair centered in each trough (leaving a 0.5-mm gap between the tip of the post and the bottom of the casting trough). Each tissue consisted of a 97-μl fibrinogen-media solution (89 μl of RPMI-B27, 5.5 μl of DMEM/F12, and 2.5 μl of 200 mg/ml bovine fibrinogen; Sigma-Aldrich) containing 5 × 10^5^ iPSC-CMs and 5 × 10^4^ supporting HS27a human bone marrow stromal cells (ATCC), which was chilled and mixed with 3 μl of cold thrombin (at 100 U/ml; Sigma-Aldrich) just before pipetting into the agarose casting troughs. The 3D-EHT mixtures were incubated for 90 min at 37°C, at which point the fibrin gels were sufficiently polymerized around the posts to be lubricated in media and transferred from the casting troughs into a 24-well plate with fresh 3D-EHT media (RPMI-B27 with penicillin/streptomycin, and 5 mg/ml aminocaproic acid; Sigma-Aldrich). 3D-EHTs were supplied with 2.5 ml/well of fresh 3D-EHT media three times per week.

In situ contractile measurements were performed at 3-weeks post casting of 3D-EHTs. To pace 3D-EHTs, post racks were transferred to a custom-built 24-well plate with carbon electrodes connected through an electrical stimulator (S88X; Astro Med Grass Stimulator) to provide biphasic field stimulation (5 V/cm for 20-ms durations) during imaging (Leonard et al., 2018). 3D-EHTs were equilibrated in Tyrode’s buffer (containing 1.8 mM Ca^2+^) preheated to 37°C and paced at 2 Hz, which was greater than the average spontaneous twitch frequency of the tissues. Videos of at least 10 contractions were recorded inside a 37 °C heated chamber using a monochrome CMOS camera (ORCA-Flash4.0). The camera lens configuration allowed for a capture rate of 66 fps with 8.3 μm/pixel resolution and a FOV of 1,536 × 400 pixels, which was sufficient to capture images of the whole 3D-EHT from rigid to flexible post. A custom Matlab program was used to threshold the images and track the centroid of the flexible post relative to the centroid of the rigid post. The twitch force profile, F_twitch_(t) = kpost × Δ_post_(t), was calculated from the bending stiffness k_post_ and deflection of the flexible post Δ_post_ at all time points (t), where k_post_ = 0.95 μN/μm was determined from beam bending theory using the dimensions of the posts and taking the Young’s modulus of PDMS to be 2.5 MPa (Sniadecki and Chen, 2007). The twitch force and twitch kinetics were calculated from the twitch force profiles using a custom Matlab program. For statistical significance testing, one-way ANOVA with a Dunnett’s correction for multiple comparisons was performed.

### Multi Electrode Array (MEA)

iPSC-CMs were plated (as above) at 100,000 cells per well directly on the electrode array of a 24 well MEA plates (CytoView MEA 24, Axion Biosystems) pre-coated with matrigel. RPMI-B27 was exchanged every two days. MEA analysis occurred at day 35 post differentiation (2 weeks post plating on MEA plates). MEA data was acquired at 37°C and 5% CO_2_ and recordings were acquired for 5 min using the Maestro MEA system (Axion Biosystems) using standard recording settings for spontaneous cardiac field potentials. Automated data analysis was focused on the 30 most stable beats within the recording period. The beat detection threshold was dependent on the individual experiment, and the FPD was manually annotated to detect the T-wave. The FPD was corrected for the beat period according to the Fridericia’s formula: FPDc = FPD/(beat period)^1/3^ (Millard et al., 2018; Asakura et al., 2015; Rast et al., 2016). Results for individual wells were calculated by averaging all of the electrodes. For each MEA experiment, 4 technical replicates per condition were averaged, and reported results represent the average of 3 distinct biological replicates. For statistical significance testing, one-way ANOVA with a Dunnett’s correction for multiple comparisons was performed.

### Fixation and immunohistochemistry

For images in Figures 2 and 5, iPSC-CMs were fixed with 4% paraformaldehyde (PFA) in PBS at room temperature for 20 min and then extracted for 5 min with 0.3% Triton X-100 and 4% PFA in PBS as previously described (Burnette et al., 2014). Cells were washed three times in 1 × PBS. Cells were blocked in 5% BSA in PBS for at least 30 minutes. Primary antibodies were diluted in 5% BSA + 0.3% Triton X-100. RBM20 antibody (ThermoFisher, Cat#PA5-58068) was used at 1:500, DDX6 (MilliporeSigma, Cat# SAB4200837), and G3BP1 (SantaCruz Biotechnology, Cat# sc-365338) antibodies were used at 1:200. For actin visualization, phalloidin-488 (ThermoFisher Scientific, Cat# A12379) in 1x PBS (15 μl of stock phalloidin per 200 μl of PBS) was used for 3 hr at room temperature. DNA was visualized via DAPI staining for 30 minutes at room temperature (final concentration 1.2 μM). Cells were kept and imaged in 1x PBS.

### Structured Illumination Microscopy

SIM microscopy was performed on a GE Healthcare DeltaVision OMX SR microscope equipped with a PLAPON 60xOPSF/1.42 NA objective with a pco.edge 4.2 sCMOS camera at room temperature. Image reconstruction was performed using GE softWoRX software. Grant S10 OD021490

### Spinning Disk Microscopy

Spinning disk microscopy was performed on a Nikon Eclipse Ti equipped with a Yokogawa CSU-W1 spinning disk head, Andor iXon LifeEMCCD camera, and 100x Plan Apo and 60x Plan Apo objectives.

### Co-localization Analysis

3-dimensional Z-stack spinning disk confocal images of RBM20 and DDX6 were separately binarized using the Allen Institute Cell Segmenter (Centrin-2 pipeline for RBM20 and Dots pipeline for DDX6, exact segmentation parameters available upon request; https://www.allencell.org/segmenter.html). Segmented channels were then max projected and co-localization was measured using the imaged-based MeasureColocalization pipeline in CellProfiler (version 4.0.4) (Carpenter et al., 2006). Measurement of co-localization reflects the Pearson’s correlation coefficient (Adler and Parmryd, 2010).

### RBM20 localization analysis

To accurately quantify RBM20 localization, analysis of SIM images was required, as confocal techniques did not provide the required resolution (see Figure 2). 3-dimensional Z-stack SIM images of RBM20 and DNA (Hoescht stain) were individually binarized using the Surfaces function in Imaris 9.7 (https://imaris.oxinst.com/). Two additional RBM20 masks were made using the nuclear mask as a fiducial marker. First, a “Nuclear RBM20 mask” was made by assigning all RBM20 within the nucleus a value of, “1”. Second, a, “Cytoplasmic RBM20 mask” was made by assigning all RBM20 outside the nuclear mask a value of, “0”. Using these masks, the intensity value of RBM20 particles was measured in the nuclear and cytoplasmic compartments, and percent localization of nucleus vs cytoplasm was calculated (e.g. percent in compartment divided by total signal). Statistical significance was calculated using a student’s T-test.

### Western blot

Protein lysates were obtained with RIPA buffer containing 1x Halt protease inhibitor cocktail (Thermo Fisher Scientific). Following clarification of the lysate by centrifugation and assessment of protein concentration by BCA assay (Thermo Fisher Scientific), samples were diluted with 3x Blue Protein Loading Dye (New England Biolabs) and boiled for 5 minutes. Twenty micrograms of protein were separated via electrophoresis using 4 – 20% Mini-PROTEAN TGX Precast Protein Gels (Bio-Rad). Proteins were transferred to PVDF membranes and blocked in TBS with 0.1% Tween-20 (TBST) and 5% Blotting Grade Buffer (BGB, Bio-Rad). The anti-RBM20 rabbit polyclonal primary antibody (Abcam, ab233147)) was diluted at 1:1000 in TBST 5% BGB and incubated overnight at 4 °C. Membranes were washed three times in TBST for 10 min at room temperature, incubated for 1 h at room temperature with goat anti-rabbit HRP-conjugated secondary antibody, and washed three times in TBST for 10 min. Chemiluminescent reaction was initiated by incubation with SuperSignal West Pico Chemiluminescent Substrate (ThermoFisher), and images were acquired using a ChemiDoc Imaging System (Bio-Rad) in “high resolution” mode. Before re-probing for the housekeeping protein GAPDH (mouse monoclonal [6C5] diluted at 1:5000; Abcam #8245) according to the same protocol but using goat anti-mouse HRP-conjugated secondary antibody, membranes were treated with Restore Plus western blot stripping buffer (ThermoFisher)for 15 minutes at room temperature, washed three times, and re-blocked. Band intensity was calculated by measuring intensity Fiji (Fiji Is Just ImageJ, https://imagej.net/Fiji) and normalized for background and GAPDH loading control (Schindelin et al., 2012).

---

**Extended Data Video 1**

WT iPSC-CMs beat spontaneously on Multi-electrode Array (MEA) plates. Black structures are electrodes. Video played at real time, 19.9 fps. Video length = 10 seconds, magnification = 20x. Brightfield microscopy.

**Extended Data Video 2**

3D-EHTs cast from iPSC-CMs spontaneously beat in culture. Flexible post (left) is moved during contraction while stiff glass post (right) remains motionless. Video played at real time, 66 fps. Video length = 5 seconds, magnification = 2x. Brightfield microscopy.

**Extended Data Video 3**

3D-EHTs cast from iPSC-CMs can be electrically paced. Tissue has been paced at 2 Hz. Video is played at real time, 66.6 fps. Video length = 3 seconds, magnification = 2x. Brightfield microscopy.

## EXTENDED DATA FIGURE LEGENDS

**Extended Data Figure 1.**
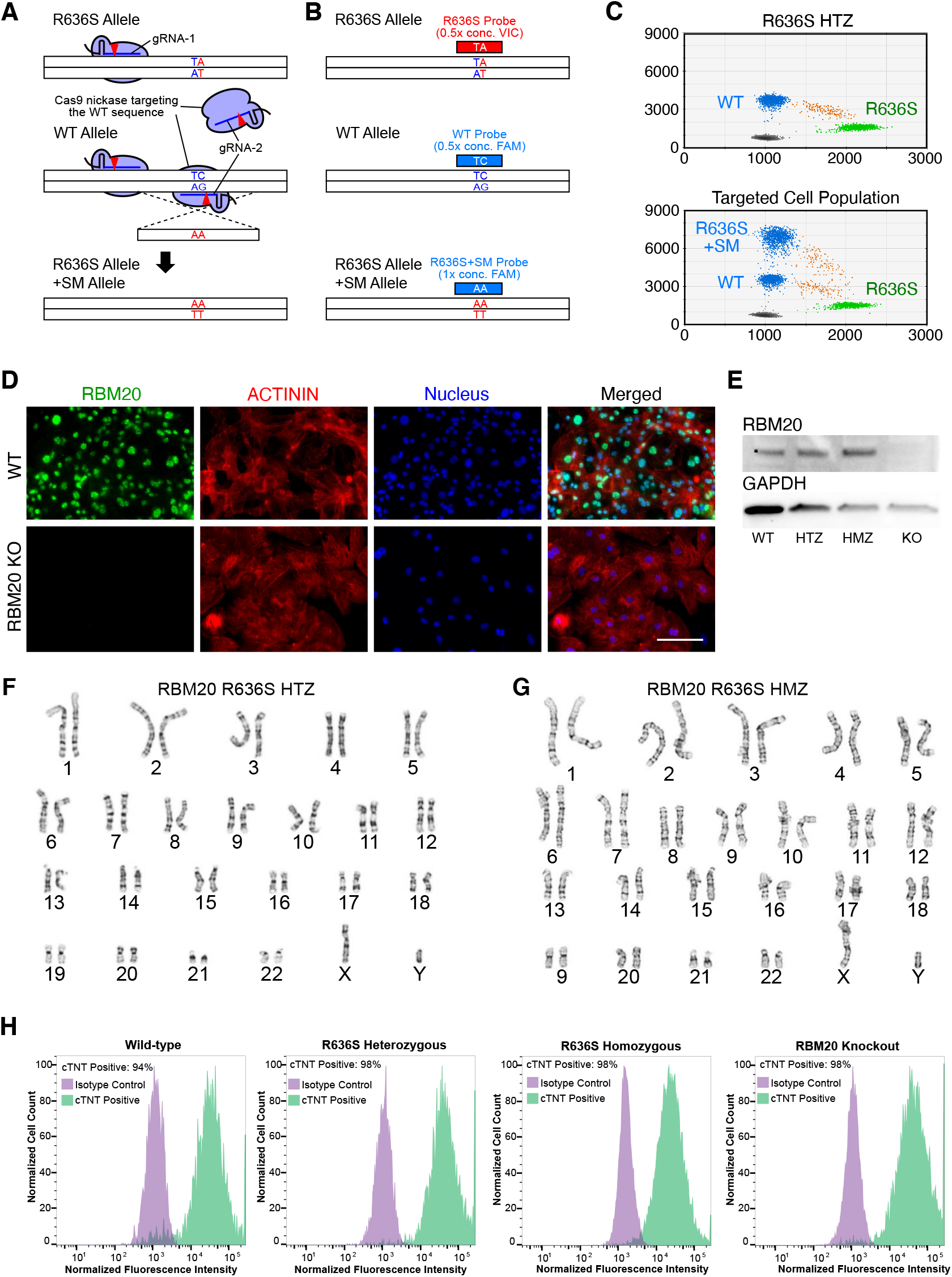
Generation of RBM20 mutant iPSC lines by genome-editing. A) Dual Cas9 nickase design to target the WT allele in R636S HTZ iPS cells to generate R636S HMZ cells. RBM20 gRNA-2 is specific to the RBM20 WT allele, so that the R636S allele could not be targeted. Recombination between the WT allele and the donor oligonucleotide with the R636S and the S635S silent mutation resulted in R636S HMZ cells that have the R636S allele and the R636S+SM allele. B) Allele specific probe design and concentration to detect the three alleles by ddPCR. To distinguish the R636S+SM allele from the other two alleles, the R636S+SM probe had two times higher concentration than the other two probes. C) ddPCR analysis to simultaneously detect the three alleles. The parental RBM20 R636S HTZ iPS cells, and a cell population targeted to generate the R636S HMZ iPS cell line were analyzed with the probes shown in B). The R636S+SM allele-positive droplets were observed in the targeted cells as FAM strongly positive population. Cell populations with a higher frequency of the R636S+SM allele and a lower frequency of the WT allele were enriched to isolate the R636S HMZ iPS cell clone. D) Immunofluorescent staining confirming depletion of RBM20 in the 8-bp Del HMZ iPSC-CMs. RBM20 (green) and ACTININ (Red) were visualized in WT and the 8-bp Del HMZ iPSC-CMs. The nuclei (blue) were stained with DAPI, and the merged images are also shown. Scale bar; 100 μm. E) Western blot showing loss of RBM20 protein in RBM20 KO iPSC-CMs. F and G) Karyotypes of R636S HTZ (E) and HMZ (F) iPS cell clones. Both lines maintained a normal male karyotype. H) Flow cytometry demonstrates RBM20 mutant iPSCs successfully differentiate into iPSC-CMs as assessed via cTnT stain following sodium lactate purification.

**Extended Data Figure 2.**
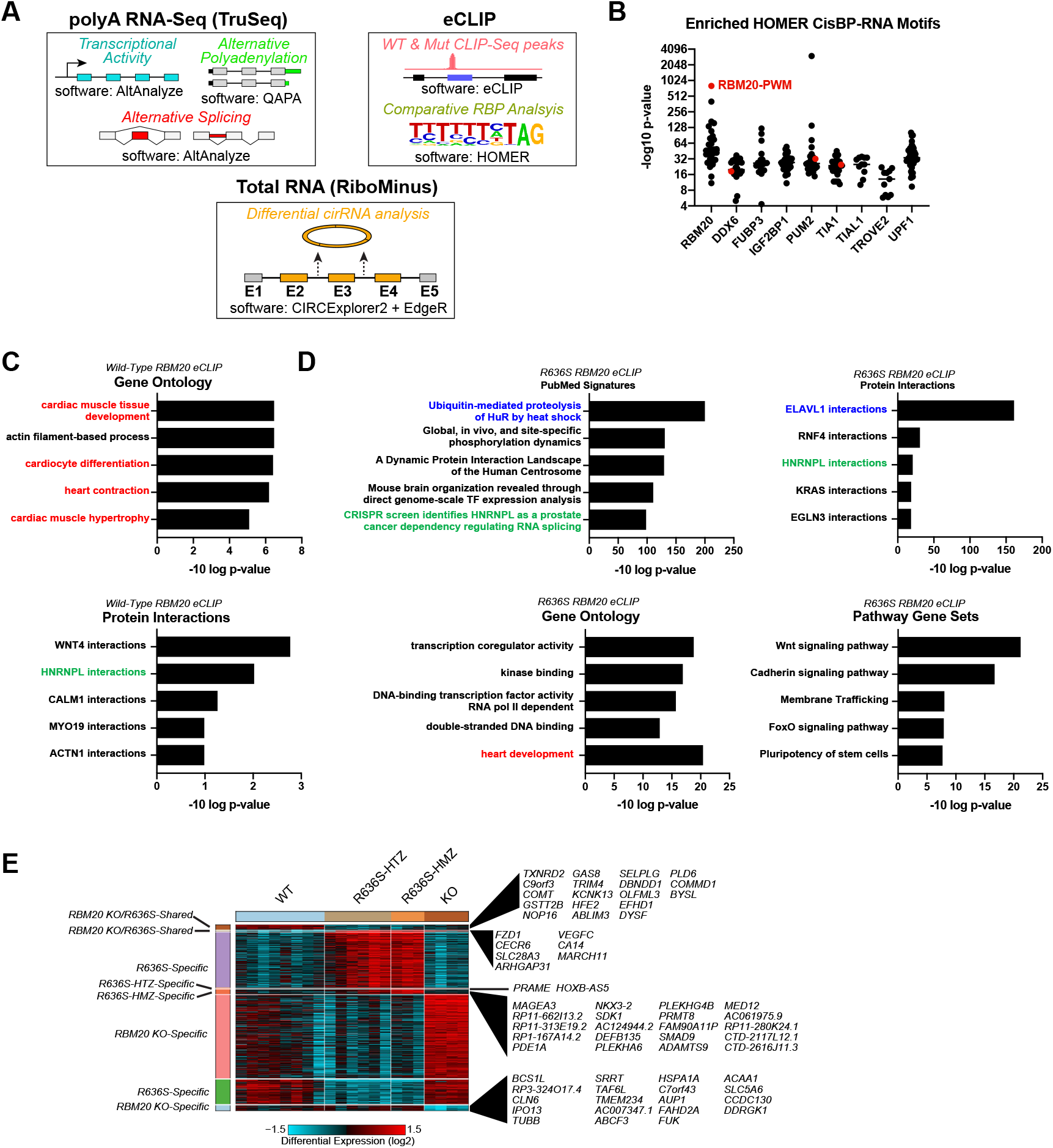
RBM20 mutant bound transcripts and functional impact. A) Graphical overview of the “omics” integrative analysis strategy and analytical approaches to identify molecular impacts of RBM20 mutation or deletion. B) Enrichment of RBM20 binding sites among RBPs with RBM20-mutant shared eCLIP peaks by HOMER using RNA recognition Elements from the CisBP-RNA database. C-D) Gene-set enrichment of analyses with the software ToppFun of genes with reproducible eCLIP peaks in wild-type (C) and R636S-HMZ (D) iPSC-CMs. E) MarkerFinder gene expression heatmap from Fig. 5, with specific genes noted for minor expression patterns.

**Extended Data Figure 3.**
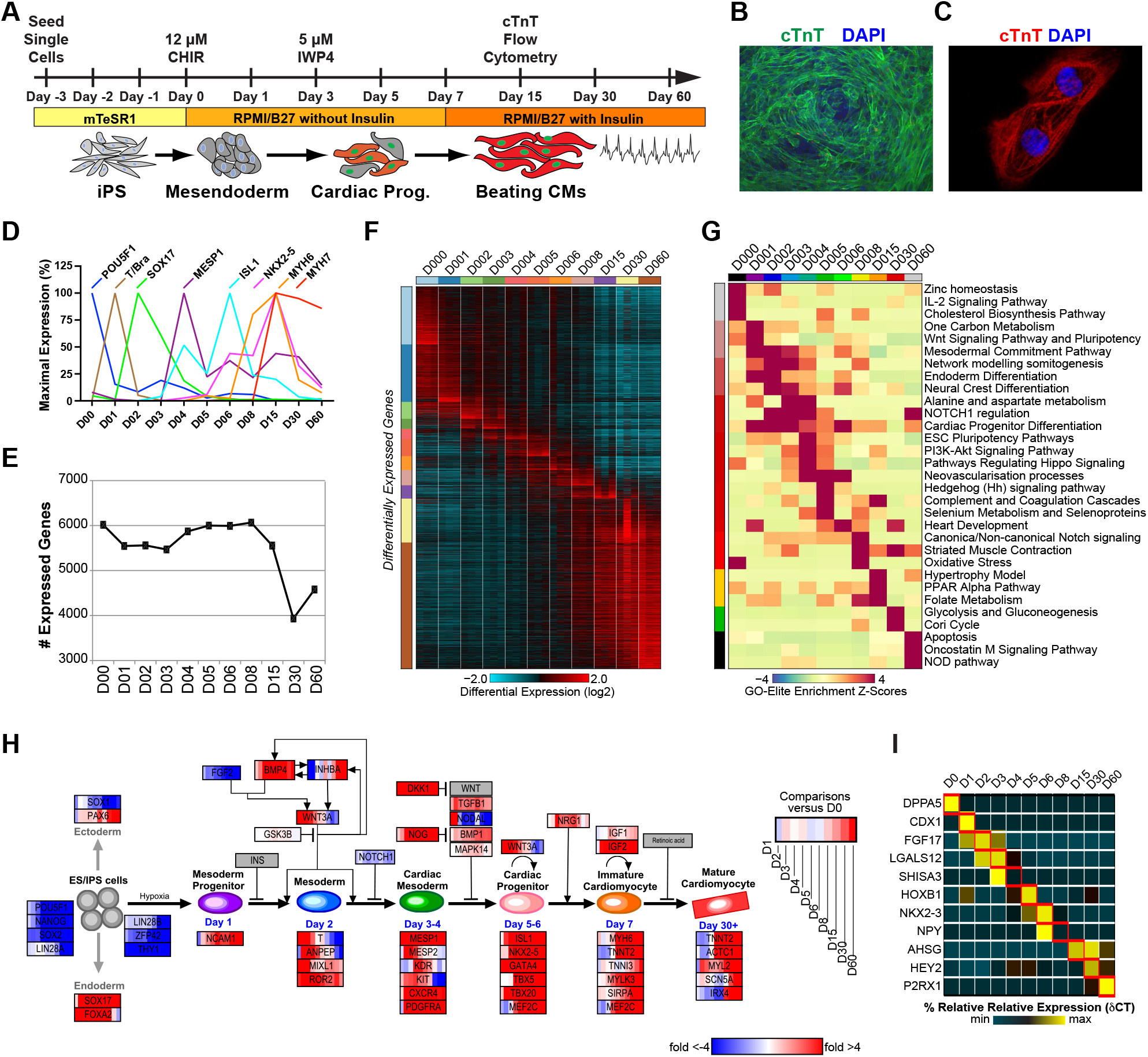
Stage-specific transcriptomic differences in cardiomyocyte differentiation. A) The GiWi protocol was used to obtain day 30 differentiated cardiomyocytes which were assessed at day 15 for negative and normal differentiated samples stained with cTnT by flow-cytometry. B-C) Differentiation efficiencies were measured by cTnT staining at (B) day 0 and (C) day 15 cells. D) TaqMan analysis shows the sequential progression from iPS cells (POU5F1) into functional CMs (NKX2-5, MYH6 & MYH7). Relative gene expression is graphed as a percentage of the maximal expression observed. E) The total number of detected genes (protein-coding and ncRNA) expressed with an RPKM > 3 and at least 50 reads/gene at each time-point of differentiation. F) Heatmap of all differentially expressed genes (fold>1.5 and empirical Bayes moderated t-test p<0.05, FDR corrected) for each timepoint for versus day 0, ordered by the software MarkerFinder. G) Heatmap of the predominant WikiPathways associated with top-200 time-point specific marker genes, based on Z-score enrichment (GO-Elite software). H) Temporal changes in gene expression (log2 fold change) across time-points of cardiac differentiation, visualized in the context of prior defined cardiac differentiation markers in PathVisio (WikiPathways:WP2406). I) Heatmap of qPCR validation results (delta CT values) of the top-predicted RNA-Seq marker genes for each time-point.

**Extended Data Figure 4.**
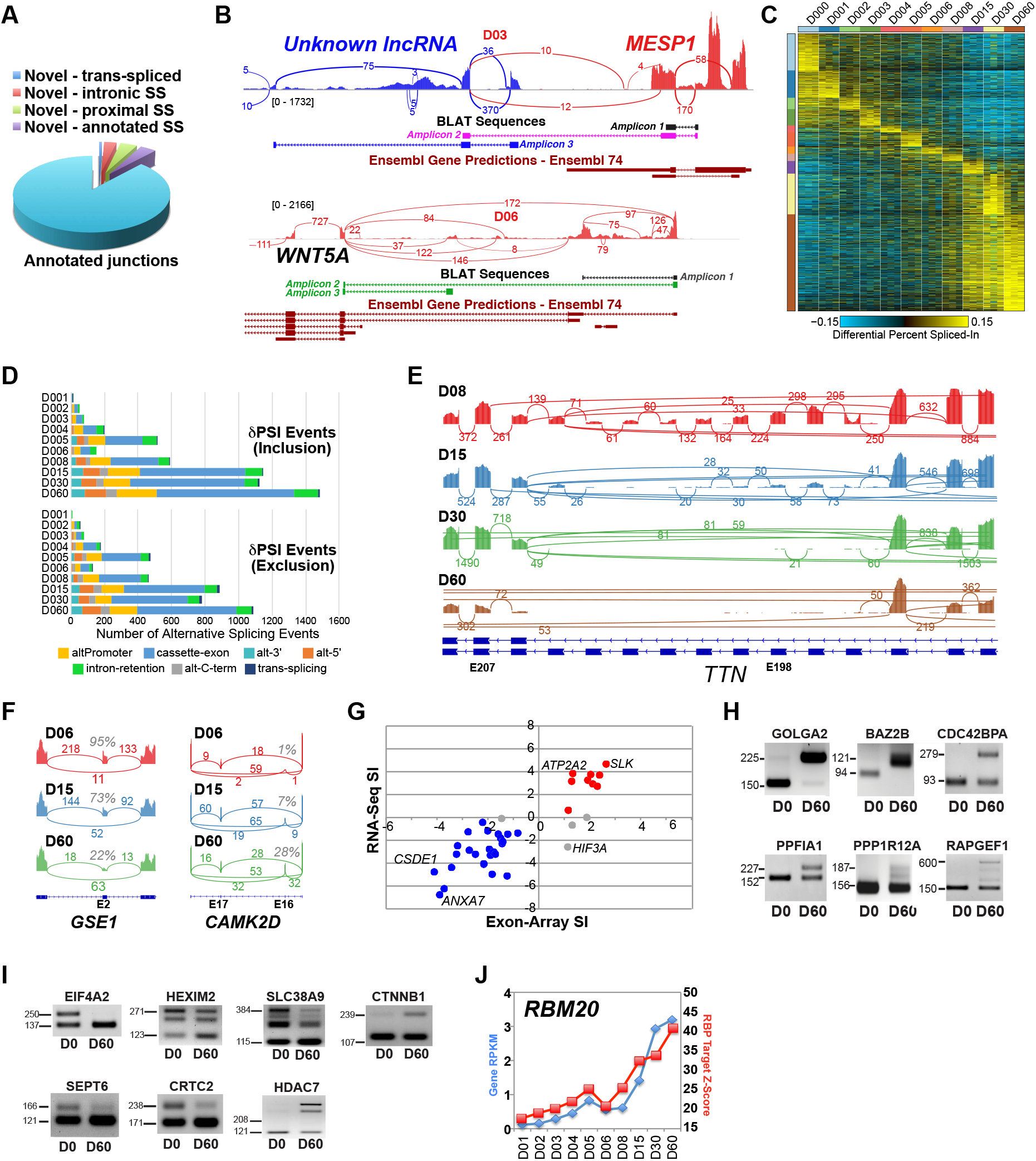
Validation of RBM20-mediated splicing during iPSC-CMs differentiation. A) Extent of expressed known and novel exon-exon junctions (at least 5 reads in 3 or more samples) observed from all cardiac differentiation RNA-Seq samples. Trans-spliced events are between two adjacent or distant genes, proximal splice-sites (SS) are those occurring within 50nt of a known splice site and intronic are those occurring >50nt away. B) Genes with novel exons were identified from AltAnalyze and verified by targeted amplification (amplicon) and Sanger sequencing. BLAT aligned sequences are derived from the consensus of forward and reverse primer amplifications. MESP1 splicing to an unknown (UNK) lncRNA on the reverse strand. One confirmed mRNA downstream for this lncRNA is shown in blue (amplicon 3), with a trans-splicing isoform produced between MESP1- and this lncRNA in purple (amplicon 2). Two novel N-terminal alternative isoforms of the gene WNT5A, with an alternatively excluded exon 2 (amplicon 2) and a novel alternative promoter regulated exon (amplicon 3) in green on the reverse strand. C) Heatmap of all differential splicing events (dPSI>0.1 and empirical Bayes moderated t-test p<0.05, FDR corrected) for each timepoint for versus day 0, ordered by the software MarkerFinder. D) Incidence of alternative splicing for exon-inclusion (top) and exon-exclusion for each iPSC-CM differentiation versus day 0, by Percent Spliced-In analysis. H) Heatmap of the top-200 unique splicing-events for all differentiation time-points. E) SashimiPlot of TTN exons spliced-out during distinct phases of iPSC-CM commitment. F) SashimiPlots illustrating the temporal regulation of novel (GSE1) and known (CAMK2D) RBM20 target exons by alternative splicing. Curved lines indicate splice-junctions with associated aligned reads. The percentage indicates the amount of relative exon-inclusion. G) Comparison of previously identified day 40 cardiac differentiation associated alternative splicing events (Salomonis et al., 2009) to similar computed exon-level splicing-index values from iPSC-CM day 30 RNA-Seq. Shown are the 41 out of 47 expressed exons that could be comparably mapped to AltAnalyze annotated exons are shown. red=positive SI fold (exon-inclusion in hESC), blue=negative SI fold (exon-inclusion in CM), grey=disagreeing. H-I) Representative RT-PCR images for novel detected iPSC-CM differentiation splicing events, denoting the expected size of each amplicon. Example novel RBM20 target splicing-events, evidenced from gene-edited iPSC-CMs, with evidence of differentiation regulation are separately called out in panel I. J) Gene expression level of RBM20 throughout differentiation (blue points), co-visualized with GO-Elite Z-score enrichment results for alternatively spliced exons in differentiation (versus day 0) relative to RBM20-mutant patient (dilated cardiomyopathy) versus control splicing events.

**Extended Data Figure 5.**
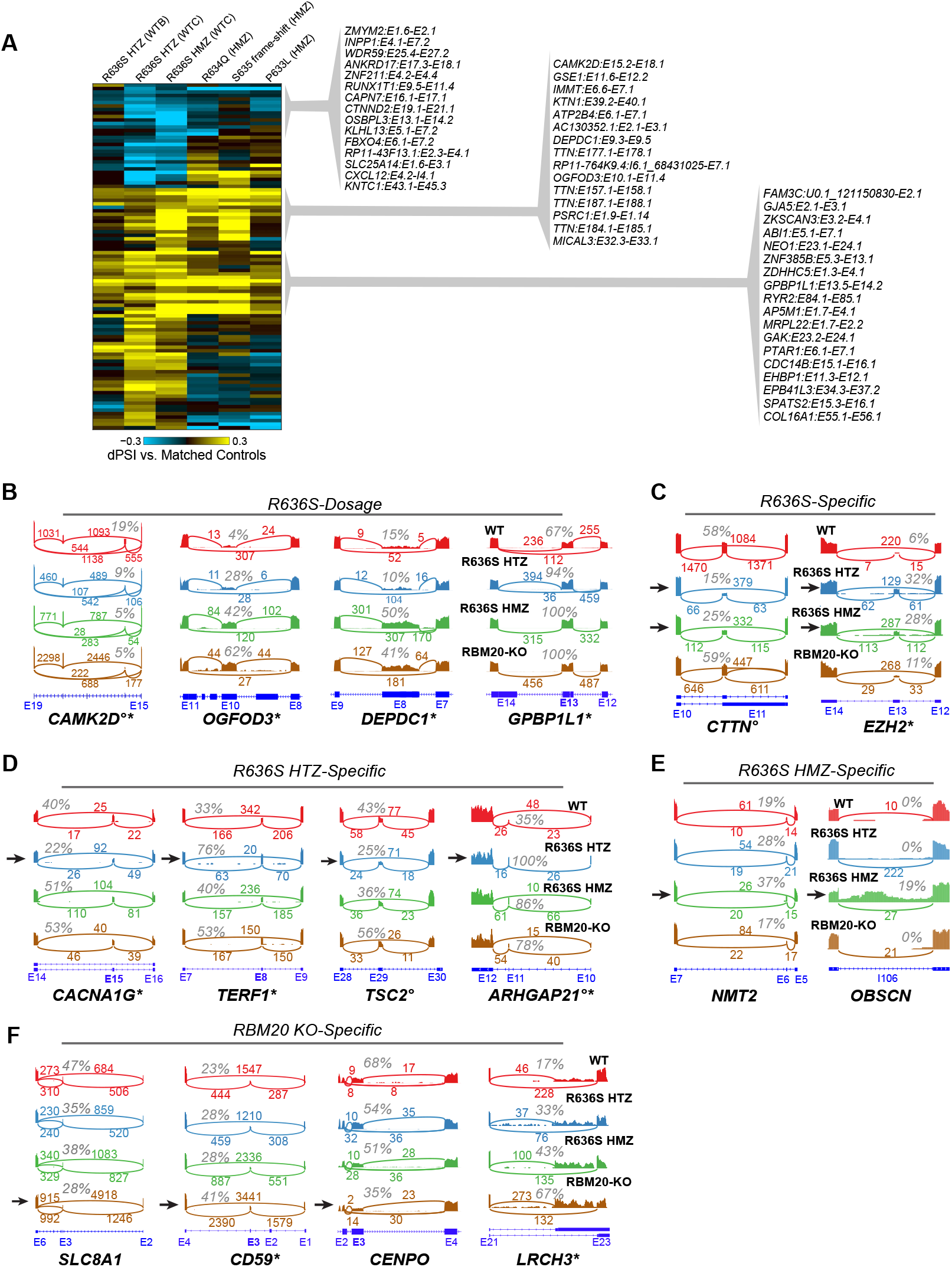
Visualization and verification of specific iPSC-CM RBM20 splicing events. A) Heatmap of shared splicing patterns observed in independently RBM20 edited iPSC-CM lines (δPSI vs. matched control lines). The R636S HTZ mutation was edited into the WTB line, whereas independently edited HMZ iPSC-CMs (including HMZ S635 frame-shift), were reanalyzed from Briganti et al. B-F) SashimiPlots of example splicing events from Fig. 4C, associated with the indicated patterns of unique or shared regulation. * = verified splicing event patterns inferred from the independently edited iPSC-CMs. ° = verified splicing event from R636S HTZ edited pig hearts.

**Extended Data Figure 6.**
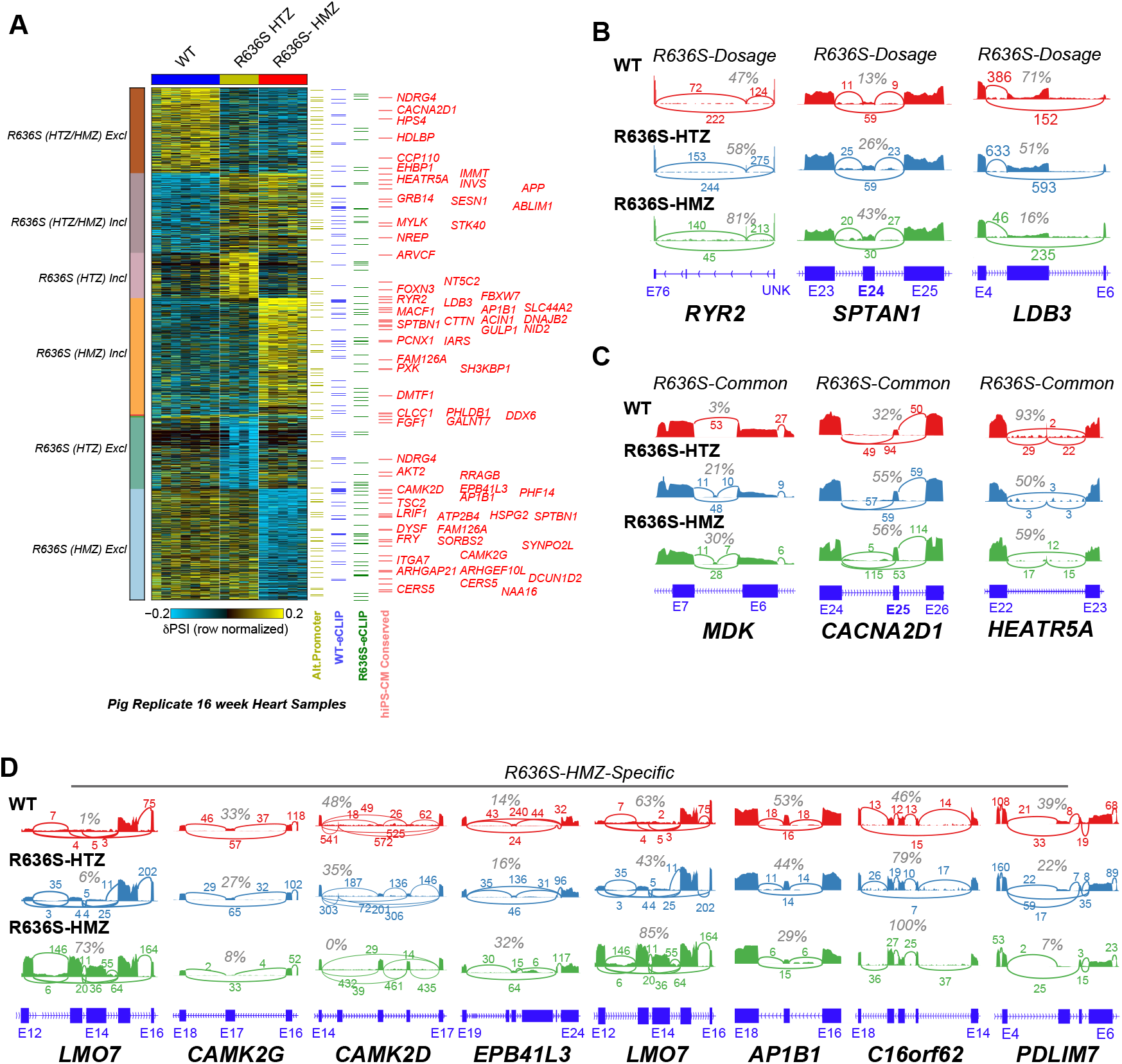
Porcine 16-week neonatal heart R636S-mutant regulated splicing events. A) Heatmap of the predominant patterns of alternative splicing in the pig heart for significantly differential events (dPSI>0.1 and empirical Bayes moderated t-test p<=0.05). See Fig. 4C for a detailed description of the overlapping eCLIP and human-conserved splicing event predictions. Genes for conserved hiPS-CM RBM20 mutant splicing events are displayed to the right. B-D) Example SashimiPlots of R636S-regulated splicing events in pigs that correspond to conserved and non-conserved events. Specifically, R636S events with observed dosage dependent splicing (B), R636S dosage-independent (C) and R636S-HMZ-specific splicing events.

**Extended Data Figure 7.**
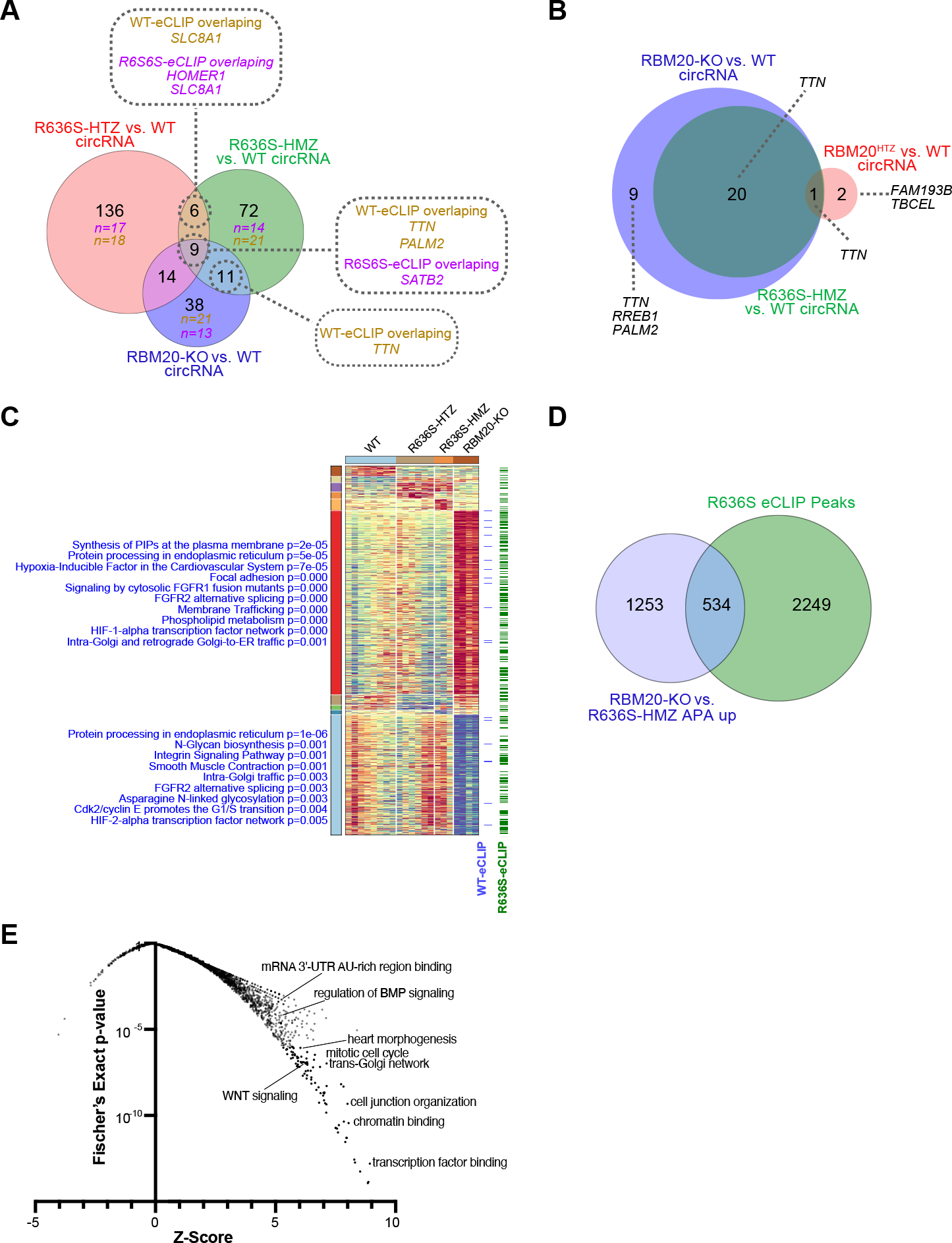
Impact of RBM20 mutation on circular RNA and alternative polyadenylation. A) Venn diagram of differentially regulated circRNAs, comparing R636S mutants or RBM20 KO to wild-type RiboMinus RNA-Seq sample groups (EdgeR p<0.1). circRNA genes with evidenced eCLIP peak genomic overlaps are denoted, based on the eCLIP source (mutant or wild-type). The number of circRNAs with eCLIP overlaps are denoted in each Venn circle (purple = R636S eCLIP, gold = WT eCLIP). B) Venn diagram of differentially expressed circRNAs that also have genomic overlapping alternative splicing events for the indicated sample-group comparisons. Such events represent coordinately regulated circRNAs and alternative-splicing events. The majority of such events are associated with a single gene (*TTN*), that are associated with multiple independent splicing events. C) Heatmap of alternative polyadenylation (APA) 3′ UTRs, organized using the software MarkerFinder into the predominant patterns of regulation. Gene-set enrichment results (GO-Elite) for each cluster are shown to the left of the heatmap (ToppFun pathway collection). Note that for each APA event, at least one reciprocal redundant APA event may also be reported (e.g., short form, long form). Any reproducible eCLIP peaks within the gene body of APA impacted genes are shown are indicated to the right of the heatmap. D) Venn diagram of genes with APA events significantly up-regulated in RBM20 KO versus R636S HMZ iPSC-CMs (relative APA ratio differences > 0.1 and empirical Bayes moderated t-test p<0.05) and genes with observed R636S eCLIP peaks. E) Gene Ontology enrichment analysis (GO-Elite) for the 534 overlapping RBM20 KO and R636S eCLIP peaks from panel D.

