## Supplemental Methods for "Gain-of-function cardiomyopathic mutations in RBM20 rewire splicing regulation and re-distribute ribonucleoprotein granules within processing bodies"

**SUPPLEMENTARY METHODS**

**ddPCR assay to detect the WT, R636S, and R636S+SM alleles**

The composition of the premixtures of allele-specific TaqMan probes and primers was 5 µM of an allele-specific FAM or VIC TaqMan MGB probe (Thermo Fisher Scientific), 18 µM of a forward primer and 18 µM of a reverse primer (Integrated DNA Technology) in water. To detect point mutagenesis, we mixed the following reagents in 0.2 ml PCR 8-tube strips: 4 µl water, 12.5 µl 2× ddPCR Supermix for probes (Bio-Rad), 1.25 µl R636S+SM FAM probe and primer premixture, 0.625 µl WT VIC probe and primer premixture, 0.625 µl R636S FAM probe and primer premixture, and 5 µl (50–150 ng) genomic DNA solution (25 µl total volume). The conditions for droplet generation, thermal cycling, and data analysis for RBM20 R636S mutagenesis with the ddPCR system were described before (ref. Miyaoka nat methods). As the R636S+SM FAM probe had a higher concentration than the WT FAM probe, the signal of the R636S+SM allele was distinguishable from that of the WT allele (Fig. S1). Cell populations with a higher frequency of the R636S+SM allele and a lower frequency of the WT allele were enriched by sib-selection until the RBM20 R636S Homo iPS cell clone was isolated.

### Oligonucleotide donor DNA used in the present study

| | |
|---|---|
| RBM20 R636S | ACAGATATGGCCCAGAAAGGCCGCGGTCTAGTAGTCCGGTGAGCCGGTCACTCTCCCCGA |
| RBM20 R636S+SM | CACAGATATGGCCCAGAAAGGCCGCGGTCAAGTAGTCCGGTGAGCCGGTCACTCTCCCCG |

The R636S point mutation and the S635S silent mutation (SM) sites are underlined and double underlined, respectively.

### gRNAs used in the present study

| | |
|---|---|
| 1 | gCCATATCTGTGAGGGAGCCAAGG |
| 2 | gAAGGCCGCGGTCTCGTAGTCCGG |

These two gRNAs were used as a dual Cas9 nickase system. RBM20 gRNA-2 specifically targeted the WT allele in R636S Het iPS cells due to the nucleotide difference at the R636S point mutation site, which is underlined. The PAM sequences are double underlined.

**Probe-primer sets used in the present study**

| | Sequence | Fluor-quencher | Final concentration |
|---|---|---|---|
| Forward primer | TGTGAAGATTCTAAATCCTGCTCCTT | | 900 nM |
| Reverse primer | AGGAGGTGAAGCTGGGAGTGT | | 900 nM |
| WT probe | CCGCGGTCTCGTAG | VIC | 125 nM |
| R636S probe | CGGTCTAGTAGTCC | FAM | 125 nM |
| R636S+SM probe | CCGCGGTCAAGTAG | FAM | 250 nM |

The R636S point mutation and the S635S silent mutation (SM) sites are underlined and double underlined, respectively.
